# 3D model of mouse embryonic pancreas and endocrine compartment using stem cell-derived mesoderm and pancreatic progenitors

**DOI:** 10.1101/2022.10.11.511696

**Authors:** Shlomit Edri, Vardit Rosenthal, Or Ginsburg, Abigail Newman Frisch, Christophe E. Pierreux, Nadav Sharon, Shulamit Levenberg

**Author notes:** Corresponding authors: Shlomit Edri and Shulamit Levenberg.

## Abstract

The developing mouse pancreas is surrounded by mesoderm compartments providing signals that induce pancreas formation. Most pancreatic organoid protocols lack this mesoderm niche and only partially capture the pancreatic cell repertoire. This work aimed to generate pancreatic aggregates by differentiating mouse embryonic stem cells (mESCs) into mesoderm progenitors (MPS) and pancreas progenitors (PPs), without using extracellular matrix substitutes. First, mESCs were differentiated into epiblast stem cells (EpiSCs) to enhance the PP differentiation rate. Next, PPs and MPs aggregated together giving rise to various pancreatic cell types, including endocrine, acinar, and ductal cells, and to endothelial cells. Single-cell RNA sequencing analysis revealed a larger endocrine population within the PP+MP aggregates, as compared to PPs alone or PPs in Matrigel aggregates. The PP+MP aggregate gene expression signatures and its endocrine population percentage closely resembled those of the endocrine population found in the mouse embryonic pancreas, which holds promise for studying pancreas development.

## Introduction

The pancreas is a gland with two main compartments, exocrine and endocrine, which are essential for digestion and glucose homeostasis, respectively. Its endocrine function is achieved through the release of glucagon and insulin, two major hormones produced by alpha and beta cells, respectively, and which are required to maintain normal blood glucose levels. The exocrine compartment is composed of acinar cells, which synthesize, store and secrete enzymes, such as trypsin, amylase and lipase that break down proteins, carbohydrates and fats. These digestive enzymes drain into the duodenum via a highly branched network of ducts within the pancreas ^1,2^.

In early stage of development, mammalian embryos undergo gastrulation in which a single layered hollow sphere of cells reorganizes into a multi-layered structure. The primary function of gastrulation is to generate an axial system, to correctly position the germ layers in relation to one another for subsequent morphogenesis and to generate the mesoderm ^3^ (Fig. S1A). In mammalian embryos, the pancreas develops from the gut tube endoderm in a region proximal to the notochord, paired dorsal aortas artery and ventral vitelline veins which all derive from the mesoderm (Fig. S1B). The mesoderm-derived cells (Fig. S1A-B) are important for generating the signals and mechanical constraints that induce the formation of the pancreatic bud and its later differentiation ^4,5^.

Pluripotent stem cells (PSCs) serve as a good model for the study of pancreas organogenesis, since they recapitulate the dynamic expression of key genes involved in this developmental process. Figure S1C provides a visual presentation of the trajectory of PSCs during in vitro differentiation, showcasing the different stages and key genes expressed in each state. The information was derived from studies investigating murine pancreas development and the differentiation of human PSCs towards pancreatic identity ^6–9^. PSCs are first differentiated to definitive endoderm (DE), which gives rise to many of the body’s internal organs. After several differentiation stages, the DE finally differentiates into multipotent pancreatic progenitors (PPs) coexpressing Pdx1 and Nkx6-1. PPs can generate the various pancreatic cell types by further differentiating into acinar, duct and endocrine progenitors ^10–12^. These, in turn, can be applied to form organoids, which are three-dimensional (3D) in vitro models of stem cells or organ-specific progenitors that have the capacity to self-organize into specific structures that display architectures and functionalities similar to in vivo tissues and organs ^13^. In addition to these interesting and promising similarities, organoids can be generated in large numbers, thereby offering the possibility to a diversity of manipulation. In the past decade, pancreatic organoids have drawn the attention of researchers due to their considerable potential in the study of pancreas development, disease modelling, drug testing and cellular therapy.

Several protocols have been developed for the generation of pancreatic organoids from human PSCs (^12,14–19^ and reviewed in ^20^). Although, mouse pancreatic organoids have also been generated from cells isolated from either adult or embryonic pancreatic tissue ^21–23^, mouse PSC-based models are less commonly used. During organoid fabrication, single cells or cell clusters are embedded in Matrigel, which provides mechanical support and serves as an extracellular matrix (ECM) substitute ^21–23^. However, the composition of Matrigel, which is a mouse tumour-derived basement membrane ECM ^24^, suffers from batch-to-batch variability, thus rendering it is suboptimal for reproducible and complete studies focusing on cell self-organization process and signals during organogenesis ^25,26^. Furthermore, this tumour-derived ECM has limited clinical translational potential ^26,27^.

Pancreatic organoids, either PSC-derived or tissue-derived organoids, are limited in their ability to fully recapitulate tissue differentiation. They also fail to reconstruct the mesoderm niche surrounding the developing pancreas. These pancreatic organoids usually lack specialized cell types, such as endothelial and mesenchymal support cells. Although there have been instances where differentiation of human PSCs into beta cells occurred along with the emergence of non-endocrine cells resembling exocrine and mesenchymal cells ^28^, such models are rare. To overcome these limitations, this work aimed to assemble a reproducible, well-defined 3D pancreatic system that recapitulates the development of the pancreas with its microenvironment. To ensure comparability of the new in vitro model with the embryonic pancreas, the initial pluripotency state of the mouse embryonic stem cells (mESCs) was assessed prior differentiation to achieve optimal pancreatic differentiation. In addition, the system integrated mesoderm to enable development of a pancreatic niche, and Matrigel usage was avoided. The work attempted to advance mESCs to epiblast stem cells (EpiSCs), then differentiate them and later form a tissue-like environment in which EpiSC-derived PPs are in contact with EpiSC-derived mesoderm progenitors (MPs). The mouse model enabled a comparative analysis by single-cell RNA sequencing (scRNA-seq) between the mouse EpiSC-derived pancreatic aggregates and the mouse embryo pancreas at various embryonic stages, which revealed the presence of various pancreatic lineages in the in vitro system. Furthermore, the single cell transcriptome in vitro – in vivo systems’ comparison enabled a deeper understanding of the limitations inherent to studying pancreas development in vitro.

## Results

### In vitro differentiation of EpiSCs to PPs passes through a rich definitive endoderm stage and results in high cell yield

mESCs differentiation usually starts when the cells are grown in the presence of either serum and leukemia inhibitory factor (LIF) ^29–31^ or 2i, a cocktail of two inhibitors of ERK and GSK3 ^32–34^. mESCs cultured in serum and LIF (referred to as SL) or in 2i are referred to as cells in a naïve or naïve ground state, respectively. Specifically, it has been shown that differentiation to pancreatic lineages started when mESCs were in a naïve state ^35,36^. The majority of works on PSCs differentiating to pancreas lineage showed advances in differentiating human PSCs ^12,36–38^. It has been demonstrated that human embryonic stem cells (ESCs) and human induced PSCs (iPSCs) exhibit greater similarities to mouse epiblast stem cells, primed PSCs, rather than to ESCs at naïve or ground states. Both human PSCs and mouse EpiSCs correspond to post-implantation epiblast and share similar morphology, signalling pathway, metabolism and epigenetic patterns ^39–42^ (Fig. 1A). This led to the postulation that advancing mESCs to EpiSCs can improve the efficacy of their differentiation to pancreatic progenitors, relying on the field advances in human PSCs differentiation. Accordingly, the response of mESCs in different states of pluripotency to pancreatic differentiation cues was examined (Fig. 1A and Methods). mESCs in naïve and in ground states and EpiSCs were subjected to a 14-day differentiation protocol based on a human PSC-to-PP protocol ^12^. At baseline, cells in the three starting conditions shared expression of the core pluripotency factors Oct4, Nanog and Sox2 (Fig. 1B). Cells grown in 2i and SL expressed higher levels of Nanog and Sox2 in comparison to EpiSCs, which also expressed the epiblast-specific gene Fgf5 that was absent in mESCs (Fig. 1B and ^43^).

**Figure 1.**
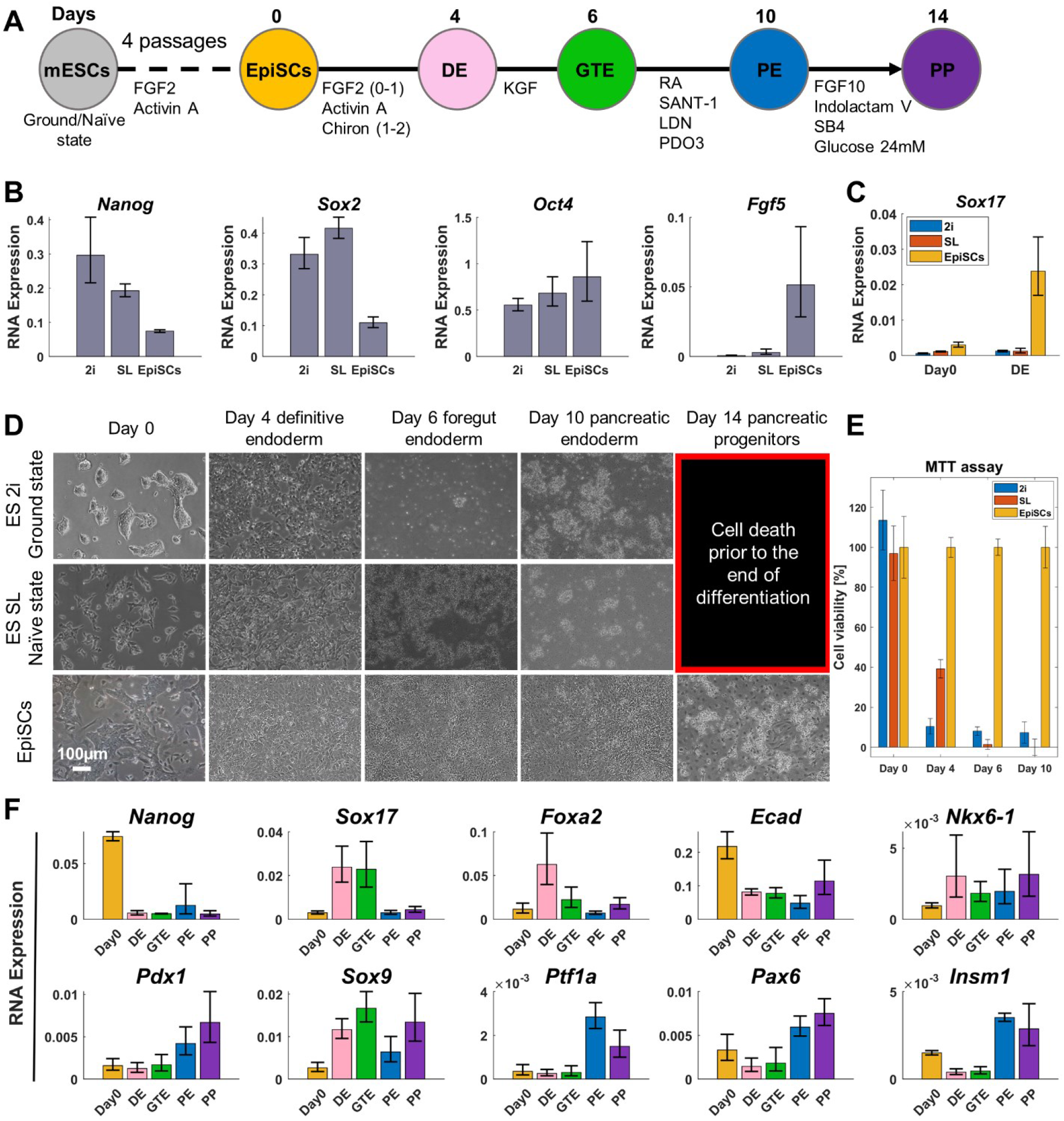
EpiSCs are a better source than ESCs for PP differentiation. **A.** In vitro differentiation course of mouse PSCs. The differentiation started either from mESCs at ground state (ES 2i), naïve state (ES SL) or EpiSCs. **B.** Relative RNA expression obtained by RT-qPCR of the three pluripotency states prior differentiation. Cells expressed the pluripotency markers Nanog, Sox2 and Oct4. Data are presented as mean±SEM, n=3 biological experiments. **C.** Sox17 expression, as measured by RT-qPCR, of cells at different initial pluripotency states during the differentiation at day 0 and day 4 referred to as definitive endoderm stage. Data are presented as mean±SEM, n=3 biological experiments. **D.** Cell morphology along the course of differentiation. **E.** Cell viability, as measured using the MTT assay, expressed as the percentage of total EpiSCs at each time point along the course of differentiation. Data are presented as mean±SEM, n=7 replicates. **F.** Relative RNA expression of pluripotency, endoderm and pancreatic gene markers during EpiSC differentiation to PPs, as measured by RT-qPCR. Data are presented as mean±SEM, n=3 biological experiments. mESCs, mouse embryonic stem cells; EpiSCs, epiblast stem cells; DE, definitive endoderm; GTE, gut tube endoderm; PE, pancreatic endoderm; PP, pancreatic progenitor.

ESCs starting their differentiation in 2i and SL growth medium, exhibited significantly lower expression of Sox17 on differentiation day 4 (DE stage), in comparison to ESCs which first transitioned to EpiSCs (Fig. 1C). Furthermore, ESCs starting their differentiation in 2i and SL growth medium, underwent cell death at around differentiation day 6 (Fig. 1D-E). In contrast, when the differentiation began after ESCs transitioned to EpiSCs, cells survived longer and a higher cell yield was obtained by the end of the differentiation period (Fig. 1D-E). Thus, it was concluded that EpiSCs are superior to naïve ESCs for endoderm and specifically pancreas differentiation, which led further investigation of EpiSC differentiation to PPs.

Along the course of EpiSC differentiation, mRNA expression (Methods and Fig. 1F) of the pluripotent markers Nanog and Ecad decreased, whilst the endodermal markers Sox17 and Foxa2 reached their highest levels at the DE and GTE stages (days 4 and 6, respectively). As the differentiation continued, Ecad expression remained constant between DE and PPs and there was a rise in the expression of pancreatic markers (Pdx1, Nkx6-1, Sox9, Ptf1a, Pax6, Insm1), mostly at PE and PP stages (days 10 and 14, respectively). At the protein level (Methods, Fig. 2 and Fig. S2), there was an elevation of pancreatic markers and a reduction in Sox17 (endoderm) and in non-pancreas endoderm lineage markers (liver, lung, stomach and intestine, see Fig. S2) at the end of differentiation. Moreover, on differentiation day 14, most of the cells coexpressed SOX9/PDX1 and over 86% of the cells coexpressed PDX1 and NKX6-1, indicating a highly efficient PP differentiation protocol (Fig. 2C and Fig. S3).

**Figure 2.**
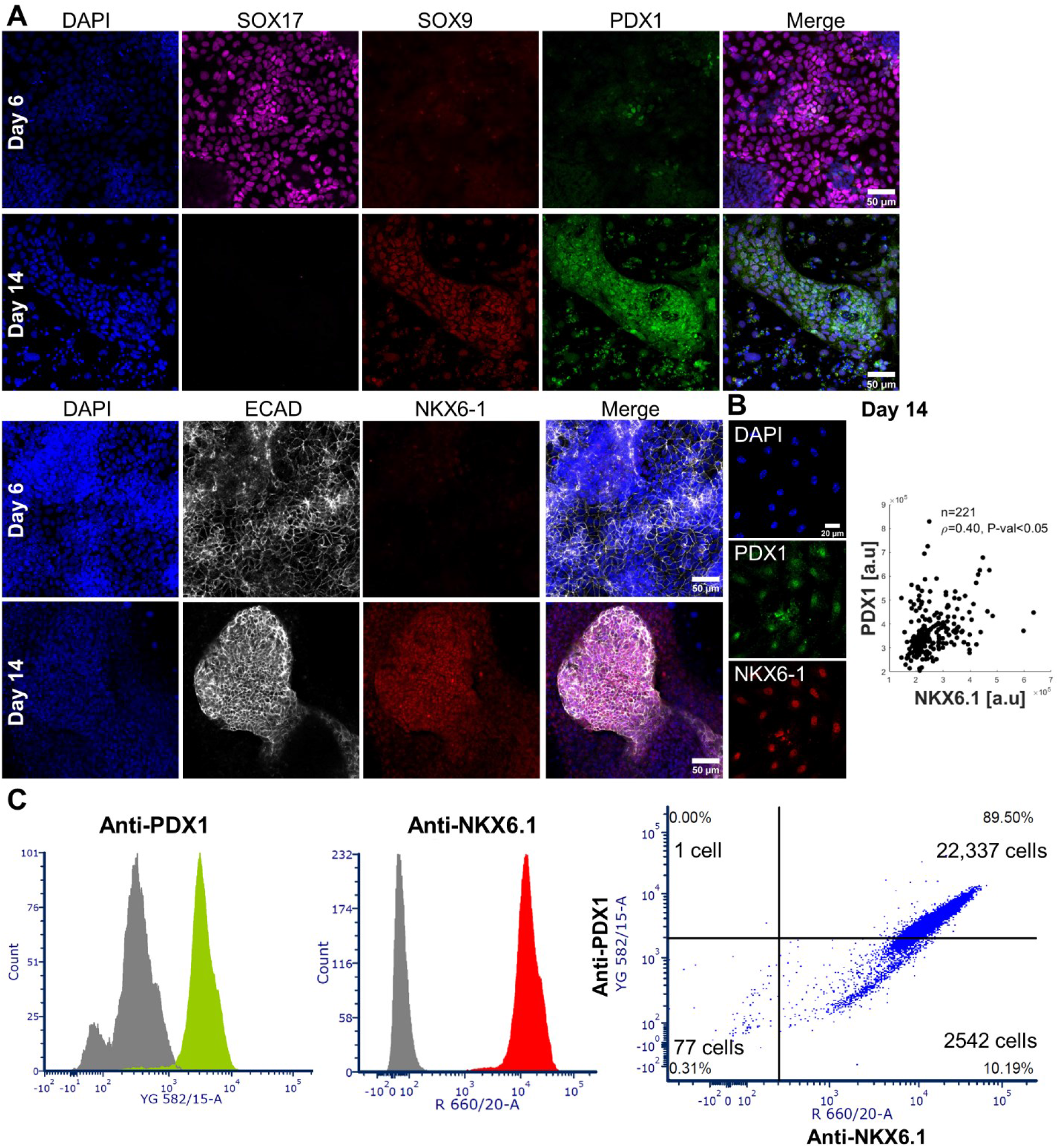
The majority of pancreatic progenitors derived from epiblast stem cells coexpressed PDX1 and NKX6-1. **A.** Confocal images captured along the course of EpiSC differentiation (day 6, i.e., the gut tube endoderm stage and day 14, i.e., the pancreatic progenitor stage at the end of the differentiation). The cells were immunofluorescently stained for endoderm marker SOX17, pancreatic markers SOX9, NKX6-1 and epithelial marker ECAD. Note that SOX17 expression decreased from day 6 to day 14, whereas pancreatic marker expression increased. **B.** Representative images of confocal images at higher magnification immunofluorescently stained for PDX1 and NKX6-1 and their fluorescence intensity quantification. The number of cells analysed is denoted by n and ρ represents Spearman’s rank coefficient with P-value <0.05. **C.** Representative flow cytometry quantification of PDX1/NKX6-1 double-positive cells on day 14 of EpiSC differentiation to PPs (n=2 biological experiments, see Figure S3).

### 3D pancreatic aggregates exhibit epithelial, acinar, endocrine and endothelial markers

When grown alone, PP aggregates did not grow in size over time (Fig. S4). We hypothesized that inductive cues from another germ layer, most probably the mesoderm, were missing hence preventing the aggregate growth. To test this hypothesis, we used EpiSC-derived mouse caudle epiblast (CE) which bear characteristics of various types of mesoderm, including allantois cells and cells resembling the notochord ^41,42^. In the present work, this CE population was tested for its ability to recapitulate the developing pancreas and its niche in the embryo. CEs, referred to here as mesoderm progenitors (MPs), were mixed with PPs in a ratio that corresponded to that observed in the mouse embryo pancreas ^44^. Each well of a 96-well plate was seeded with a 1000-cell mixture (Fig 3A), and cells aggregated to form a round structure within 3 days. Culture continued for another 4 days on an orbital shaker (Fig 3B).

**Figure 3.**
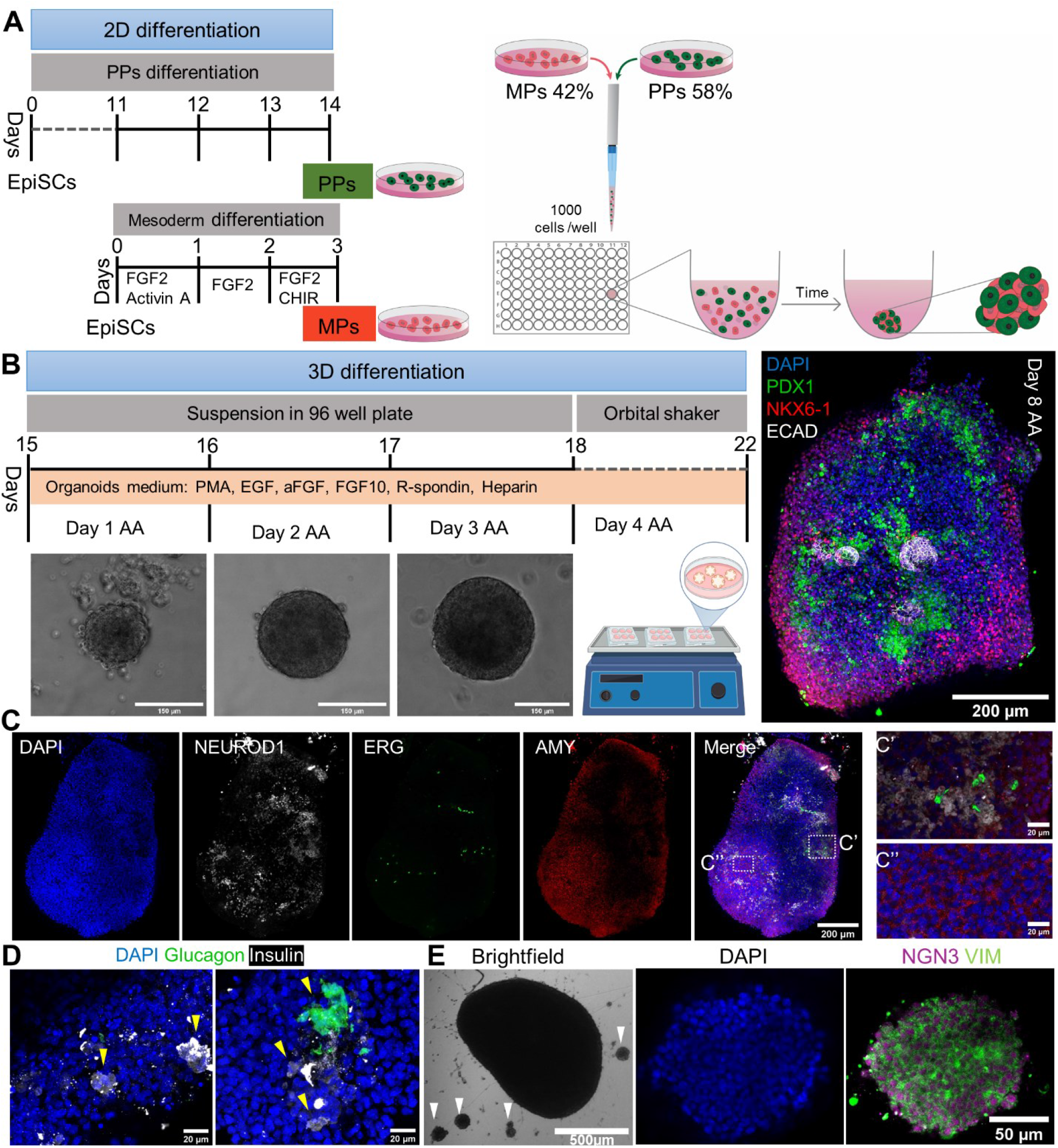
3D co-culture of pancreatic and mesoderm progenitors recapitulates the developing pancreas in the embryo. **A-B.** A schematic illustration of pancreatic progenitor (PP) + mesoderm progenitor (MP) aggregate preparation. Prior to PP and MP aggregation, 2D differentiation courses of 14 days and 3 days were carried out to obtain PPs and MPs, respectively. **A.** On day 14, 1000 PPs and MPs were mixed at a proportion of 58% and 42% and were plated to each well of a U-bottom 96-well plate. **B.** Brightfield images of the aggregates on days 1, 2 and 3 after aggregation (AA). On day 4 AA, the aggregates were moved to a 6-well plate and placed on an orbital shaker for an additional 4 days. Confocal image of the day 8 AA aggregate immunofluorescently stained for ECAD, NKX6-1 and PDX1 (for higher magnification images see Fig. S5). **C-D.** Confocal images of day 8 AA aggregates immunofluorescently stained for endocrine marker NEUROD1, acinar marker AMY (amylase), endothelial marker ERG (C, C’, C’’), and glucagon and insulin, which mark alpha and beta cells, respectively (D). Nuclear staining with DAPI is in blue. Yellow arrows in D indicate regions of cells expressing insulin or glucagon. **E.** Brightfield image of a pancreatic aggregate on day 8 AA. The white arrows in the image show small aggregates that are approximately 100-150 µm in diameter, located around the big aggregate. A representative confocal image of a small aggregate immunofluorescently stained for vimentin (Vim), NGN3 and nuclei (DAPI).

Co-culture of PPs with MPs gave rise to pancreatic aggregates expressing various pancreas-specific genes along with mesenchymal and endothelial genes. On day 8 after aggregation (AA) the PP+MP aggregates expressed the pancreatic markers PDX1 and NKX6-1, contained epithelial regions marked by ECAD (Fig. 3B), endocrine cells expressing NEUROD1 and NGN3 (Neurog3), regions of acinar cells exhibiting AMY (amylase) expression, duct cells expressing SOX9 and weak expression of PDX1 and endothelial cells marked by ERG and CD31 expression (Fig S1C and Fig. 3 and Fig. S5-S6). Furthermore, in several regions in the PP+MP aggregates, endocrine cells had differentiated into beta and alpha cells expressing insulin and glucagon, respectively (Fig. 3D, Fig. S6C).

Spatial proximity was noted between cells expressing the endothelial marker ERG and cells expressing the endocrine marker NEUROD1 (Fig. S5). Specifically, cells in the outer region of the aggregate expressed AMY, whilst cells deeper in the tissue expressed less AMY and more NEUROD1 and ERG. This alignment between endothelial and endocrine cells (CD31 and NEUROD1 in Fig. S5, ERG and NEUROD1 in Fig. S6) mimics the preferential endothelial-endocrine cell organization found in the pancreas. Previous studies demonstrated the importance of an endothelium for pancreas morphogenesis and differentiation and for induction of endocrine differentiation in the embryo ^5,45–48^.

During the 3D culture, cells at and around the aggregate boundary had an elongated morphology, and formed several small 50-100 μm aggregates, suggesting that the cells were mesenchymal with migratory characteristics (Fig. 3E). Immunostaining of the small aggregates showed that most of the cells expressed the epithelial-mesenchymal transition (EMT) marker vimentin (Vim) and the transient endocrine marker NGN3 (Fig. 3E), similar to the profile of differentiating endocrine progenitors in the embryo ^49–53^.

### Single-cell RNA sequencing analysis revealed abundant representation of mesenchymal and endocrine populations in the PP+MP aggregates

To gain a comprehensive understanding of the PP+MP aggregates and uncover the different cell populations within them, the aggregates were subjected to single-cell RNA-sequencing (scRNA-seq). Additionally, to assess the benefits provided by the mesodermal compartment within the pancreatic aggregates, a comparative analysis was conducted at the single-cell transcriptomic level between PP+MP aggregates and aggregates consisting solely of PP cells. A 3D culture of PPs mixed with Matrigel was also included in the comparison, since Matrigel is commonly employed in the construction of pancreatic organoids.

Following quality control filtering (as described in the Methods section), scRNA-seq of the 3D in vitro pancreatic aggregates yielded a total of 5056 cells from the PP_org samples, 3485 cells from the PP+Matrigel samples, 1456 cells from the PP+MP_B1 samples, and 3602 cells from the PP+MP_B2 samples. A graph-based clustering algorithm revealed 12 clusters, which were visualized using the Uniform Manifold Approximation and Projection (UMAP) dimensionality reduction technique (Fig. 4A). Looking at the sample composition of the clusters (Fig. 4C) and at the genes of interest (Fig. 4B, D-F), revealed that clusters 0 and 1 were mainly comprised of PP_org and PP+Matrigel samples (Fig. 4C) and expressed laminin protein-encoding genes and FGF genes. Clusters 3 and 7, which were mainly composed of PP+MP samples, and cluster 10, which was mainly composed of PP+Matrigel cells, had mesenchymal identity and expressed ECM markers. All the clusters and predominantly clusters 0, 1, 3, 6, 7 and 8 showed EMT markers. Cluster 2, which was mainly composed of PP_org cells, and clusters 4 and 5, which were mainly composed of PP+MP samples, expressed cyclin genes. Cluster 8, which was exclusively comprised of the PP+MP samples, contained a cell population exhibiting a pancreatic progenitor signature (Onecut1, Onecut2, Pax6) and together with cluster 6, which was mainly composed of PP+MP samples, predominantly held endocrine progenitors (Sox4, Chga, Chgb, Neurod1) and included various endocrine cell types (alpha, beta and delta cells). Although some cells exhibited expression of genes found in alpha and beta cells (Fig. 4F and Table S1), expression of insulin (Ins1 and Ins2, beta cells) and glucagon (Gcg, alpha cells) was not detected in the scRNA-seq dataset. Only somatostatin (Sst, see Table S1), which is secreted by pancreatic delta cells, was detected in the dataset. Clusters 9 and 10 were mainly composed of PP+Matrigel cells and expressed proliferation genes and ductal markers (Krt19, Muc1, Cdh1 and Itga6), respectively. Cluster 11, which was exclusively composed of PP+MP samples, had endothelial identity (Pecam1 and Cdh5). Several cells across the clusters expressed exocrine progenitor markers (Ptf1a, Bmp7). Most of the cells expressed Vim (Fig. 4F), suggesting migratory and motility capacities. Many cells expressed Cdh2, also known as N-cadherin. A study that explored the expression of N-cadherin in normal human tissues suggested that N-cadherin expression is closely associated with hormone-producing cells in the pancreas but is not expressed in exocrine cells ^54^. Indeed, higher expression of Cdh2 was measured in the endocrine clusters 6 and 8 and lower expression was measured in exocrine cluster 10 (Figure 4F).

**Figure 4.**
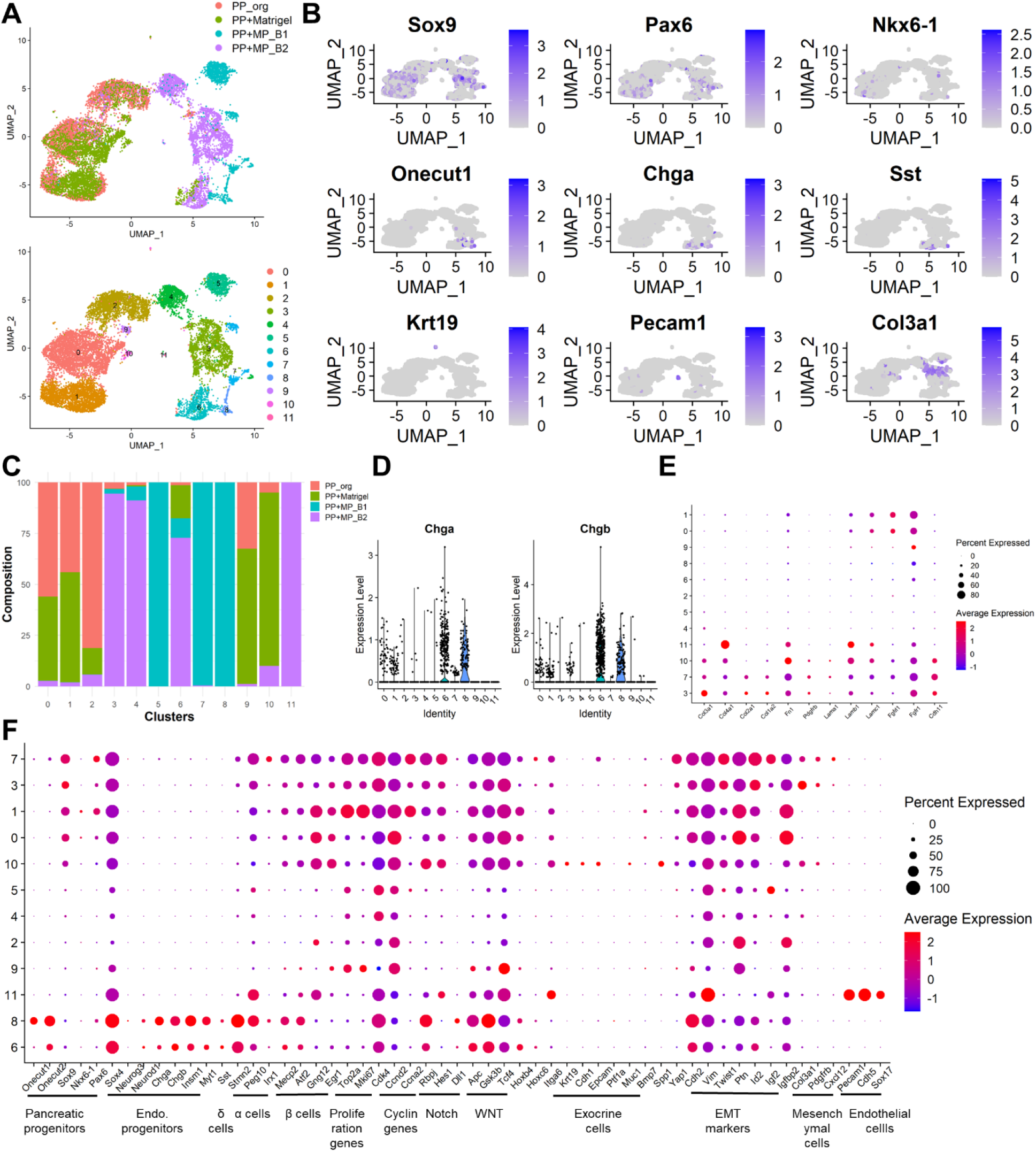
PP+MP aggregates show mesenchymal identity and contain endocrine populations. **A.** Upper graph shows dimensionality reduction UMAP plot of pancreatic aggregates coloured according to sample identity: PP aggregates (PP_org), 3D culture of PPs mixed with Matrigel (PP+Matrigel) and two biological replicates of PPs with MPs aggregates (PP+MP_B1, PP+MP_B2). At the bottom, UMAP plot of pancreatic aggregates grouped using the Seurat graph-based clustering method. The clustering algorithm (Methods) identified 12 clusters. **B.** Expression of select genes projected on the UMAP plot. Colour intensity indicates level of expression. **C.** Bar plot showing sample composition of each cluster. **D.** Violin plot visualising distribution of endocrine genes Chga and Chgb expression in each cluster. **E-F.** Expression dot plot of select genes across clusters. Clusters are ordered hierarchically based on the expression of select genes. The size of the dot corresponds to the percentage of cells expressing a gene in each cluster. The colour represents the average expression level. **E.** Expression dot plot of ECM genes, mesenchymal genes and fibroblast growth factor receptors across clusters. **F.** Expression dot plot of select genes grouped by signalling pathway and cell identity categories (see Table S1). UMAP, uniform manifold approximation and projection. Endo. progenitors, endocrine progenitors.

In general, the PP aggregates demonstrated prominent expression of Sox9, cyclin genes, laminin genes and EMT markers. The PP+Matrigel sample primarily expressed Sox9, cyclin genes, ECM genes, EMT markers and ductal markers. The PP+MP clusters predominantly exhibited expression of EMT markers, with certain populations showing a dominant endocrine and mesenchymal identity associated with cyclin genes, similar to those observed during embryonic development (^8,55–57^ and Table S1).

Focus was then placed on the pathways related to the endocrine lineage, including the expression of genes related to Notch and Wnt signaling pathways, as well as Yap1, which is part of the hippo signaling pathway. These pathways play a significant role in determining endocrine versus ductal cell lineage ^58,59^. Clusters 6-8 had low expression of Notch markers and high expression of Wnt markers, suggesting their endocrine identity.

Hox gene expression profiles provide spatiotemporal information of the emerging embryonic axial tissues (^60^ and reviewed in ^61^). Exploring their expression in the aggregate might help anchor it to a developmental stage and region in relation to the embryo. Hoxb4 was mainly expressed in PP_org and PP+Matrigel clusters 0, 1 and 10, and PP+MP clusters 6 and 8, whereas Hoxc6 was mainly expressed in cluster 7, suggesting that cluster 7 defines a more advanced developmental stage and more posterior embryonic region as compared to the other clusters. Furthermore, Hoxb4 takes part in limiting the endoderm response to signals from the notochord during the formation of the dorsal pancreatic bud ^62^. This early stage of the pancreas might correlate with the progenitor population residing in cluster 8. Hoxc6 is expressed exclusively in the mesoderm of the developing pancreas and is needed for endocrine cell differentiation ^62,63^, suggesting a mesodermal state for cluster 7, which exclusively consisted of PP+MP aggregate cells.

### The pancreatic aggregates captured diverse pancreatic cell types found in the mouse embryonic pancreas

To further characterize the co-culture system and evaluate its limitations, scRNA-seq data obtained from the pancreatic aggregates was compared to that of mouse embryo pancreases at E12, E14 and E17 (^8^ and see Methods).

Integration of the 3D in vitro and embryonic datasets yielded a UMAP with 20 cell clusters (Fig. 5A-B). Apart from identifying the different cell types, such as mesenchymal (Col3a1), ductal (Krt19) and acinar (Cpa2) cells, the key gene expression patterns (Fig. 5C) reflected a transition from a pancreatic progenitor population (cluster 11: Pdx1, Sox9 and Nkx6-1) to an endocrine progenitor population (cluster 13: Neurog3), which was later committed to an endocrine lineage (cluster 5: Chga, Neurod1, Ins1 and Gcg; Fig. 6C). Interestingly, high levels of Vim expression were recorded in all cell populations, excluding the endocrine population, consistent with the observation that Vim is lost just after the hormones are turned on during endocrine differentiation ^49^. Exploring the marker gene profiles of each cluster (Methods, Fig. 5C and Table S2) enabled determination of the cell identity of each cluster, which is summarized on the UMAP plot in Figure 5D. A clear pancreatic identity was not assignable to cluster 2, which marked by genes related to differentiating stem cells and amplifying cells, hence this cluster was called proliferating cells.

**Figure 5.**
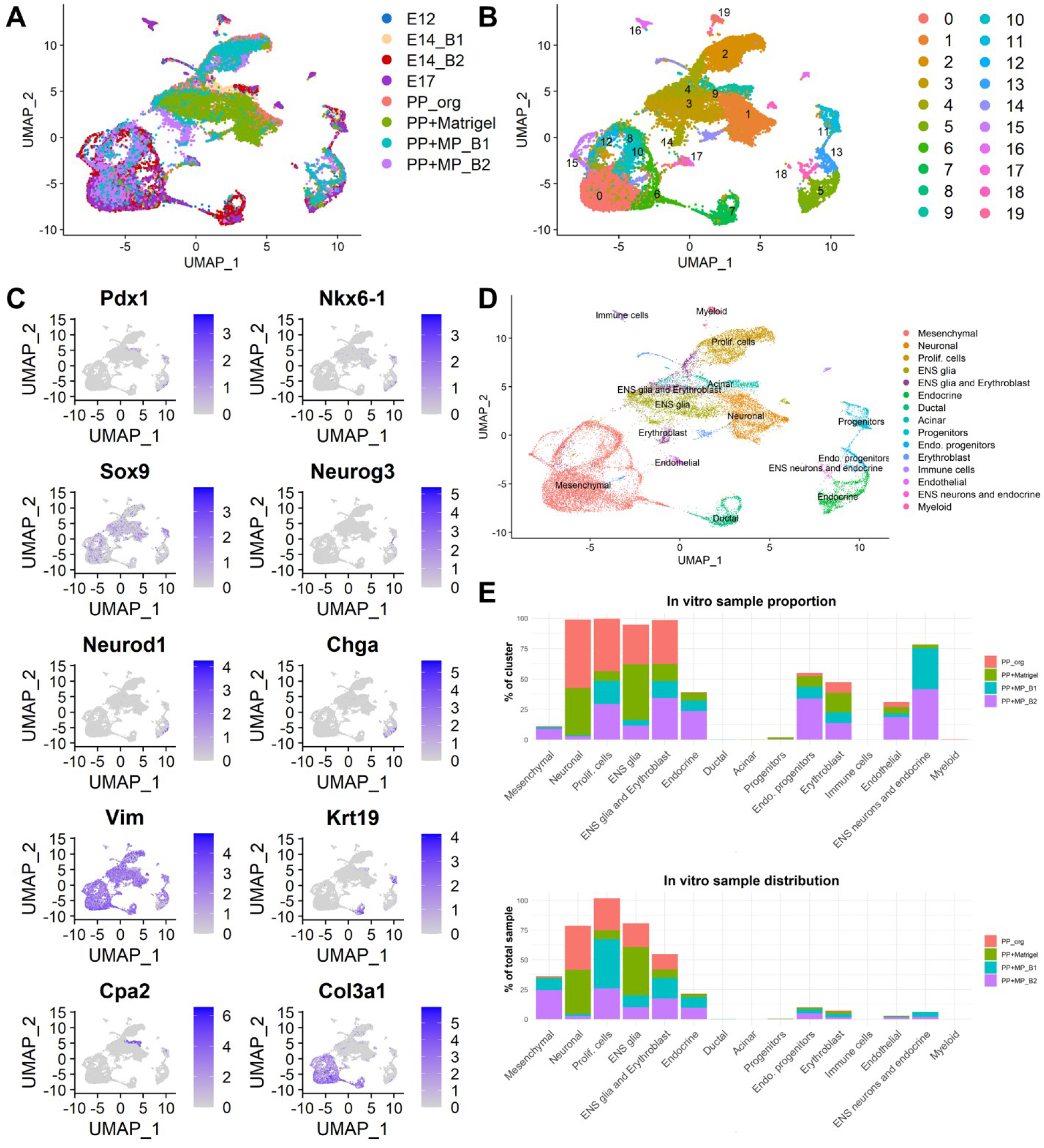
Comparison of single-cell RNA-seq of pancreatic aggregates and mouse embryo pancreases reveals cell population overlap. **A.** UMAP visualization of single-cell RNA sequencing of pancreatic aggregates: PP aggregates (PP_org, 5056 cells), 3D culture of PPs mixed with Matrigel (PP+Matrigel, 3485 cells) and two biological replicates of PPs with MPs aggregates: PP+MP_B1 (1456 cells) and PP+MP_B2 (3602 cells), and of the mouse embryonic pancreas at E12 (4412 cells), two batches at E14 (E14_B1 (3495 cells) and E14_B2 (4309 cells)) and E17 (2241 cells). Datasets were integrated using the Seurat integration algorithm. **B.** Seurat graph-based clustering revealed 20 clusters. **C.** Gene expression of select genes projected on the UMAP plot. Colour intensity indicates level of expression. **D.** A UMAP plot showing the assignment of cell type to clusters. Clusters were identified based on the expression of known markers of different pancreatic cell types as in C and as detailed in Table S2. **E.** Upper graph shows the percentage of the in vitro cells in each cell group and the bottom graph shows the distribution of the in vitro cells amongst the different cell groups. Prolif. cells, proliferating cells; Endo. progenitors, endocrine progenitors; ENS, enteric nervous system.

**Figure 6.**
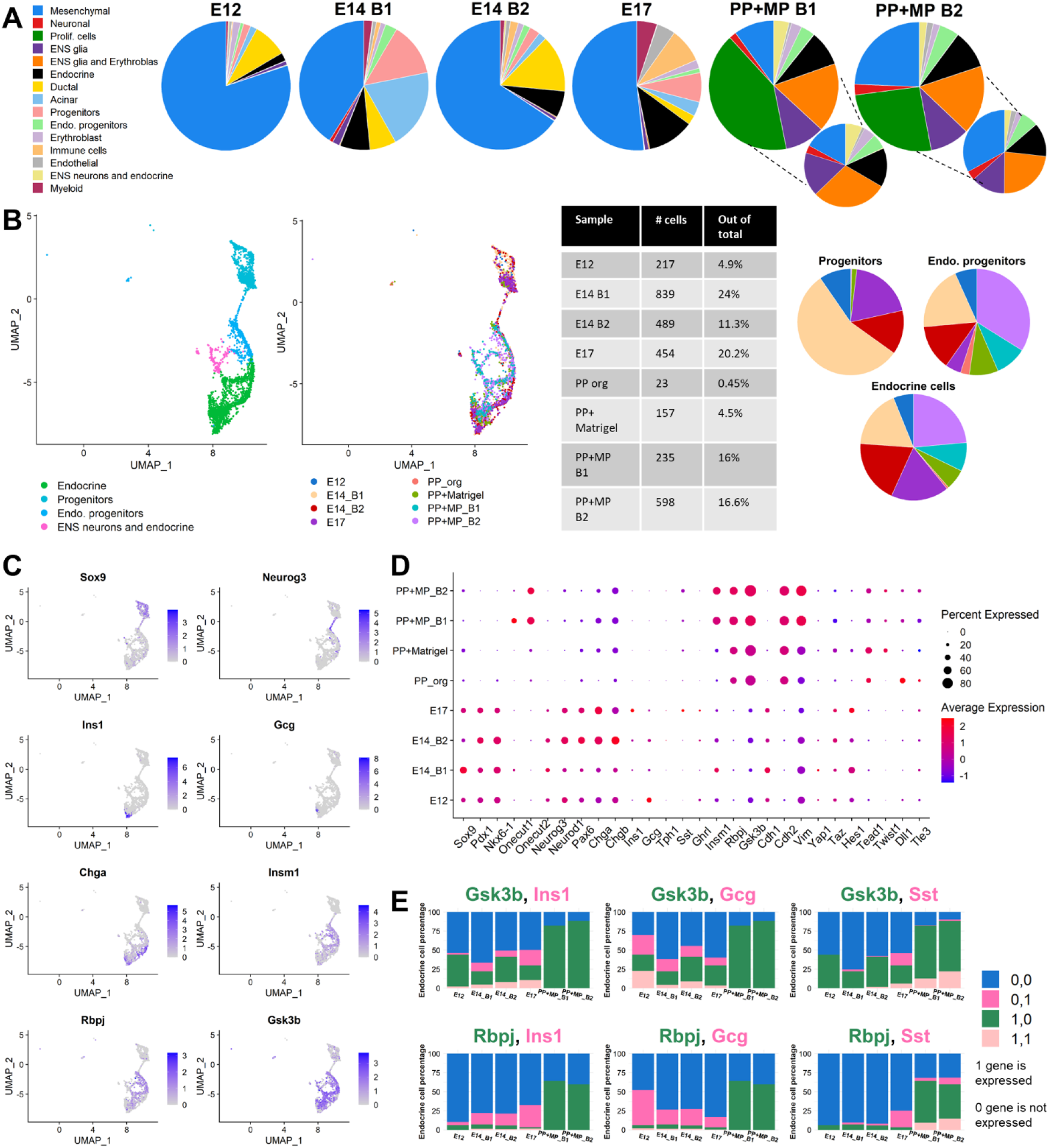
Most of the in vitro to in vivo similarity is in the endocrine compartment. **A.** Pie plots showing the proportion of the different cell types in the mouse embryo at E12, E14 and E17 and in the 3D in vitro pancreatic samples. The lower pie plots show the proportion of the different cell types in the PP+MP samples upon exclusion of the proliferating cell cluster. **B.** UMAP of four cell types: progenitors, endocrine progenitors, endocrine and enteric nervous system (ENS) neurons and endocrine cells, coloured by cell types (left) and samples (middle). Table summarises the percentage of endocrine cells in each sample and pie plots (right) show the sample distribution in the clusters of the progenitors, endocrine progenitors and endocrine cells. The sample colour legend is presented in middle UMAP plot. **C.** Gene expression of select genes projected on the UMAP plot. Colour intensity indicates level of expression. **D.** Expression dotplot of select genes in each sample. The size of the dot corresponds to the percentage of cells expressing the gene in each sample. The colour represents the average expression level. **E.** Composition of endocrine cells (clusters: endo. progenitors, endocrine and ENS neurons and endocrine) with double expression of either Gsk3b or Rbpj with the hormones Ins1, Gcg or Sst. 0,0 indicates neither expression of the 2 genes in a cell, 0,1 indicates no expression of Gsk3b or Rbpj and expression of the hormones (Ins1, Gcg or Sst) in a cell, 1,0 indicates expression of Gsk3b or Rbpj and no expression of the hormones (Ins1, Gcg or Sst) in a cell and 1,1 indicates expression of both genes in a cell.

Notably, the PP+MP aggregates displayed a distinct profile with a more abundant representation of endocrine cells compared to the other in vitro samples (Fig. 5E). The PP+MP aggregates displayed a diverse composition of cell types, as observed in the embryonic samples. Primarily, these aggregates contained mesenchymal, endocrine and proliferating cells along with erythroblasts and enteric nervous system (ENS) glia (Fig. 5E). In contrast, both PP and PP+Matrigel aggregates exhibited fewer endocrine or mesenchymal cell types, with a predominant presence of neuronal and erythroblast identities. Furthermore, 1.4% of the PP+MP cells were endothelial, which closely matched the proportions in the E14 embryo (batch 1 with 0.92% and batch 2 with 1.04%, Table S4). In contrast, the other in vitro samples demonstrated a lower endothelial cell percentage (Table S4), which suggests that inclusion of mesenchymal cells in the PP+MP aggregates was important for the generation of endothelial cells closely resembling the physiological proportion during pancreas development.

In the embryonic dataset, the mesenchymal population emerged as the predominant cell population, whereas in the PP+MP aggregate samples, cluster 2, defined as a population of proliferating cells, were dominant (Fig. 6A). Neuronal and ENS glia cell types were the dominant populations within both PP aggregates and PP+Matrigel samples (Fig. S7A). Approximately 16% of the PP+MP aggregates had an endocrine identity (Fig. 6B), which is similar to the endocrine percentage in the embryonic samples at E14 (batch 1 with 24% and batch 2 with 11%) and E17 (20%, Fig. 6B). Upon exclusion of cluster 2 from the PP+MP aggregate datasets, the mesenchymal population became prominent and a significant endocrine compartment, comprising endocrine progenitors, endocrine cells and ENS neurons, was observed (Fig. 6A).

In addition to the similarity between the endocrine proportion of the PP+MP aggregates versus the in vivo system, the endocrine differentiation path was also comparable. The in vitro cells nicely projected onto the endocrine differentiation trajectory obtained in the UMAP plot (Fig.5C-D and Fig. 6B-C), from progenitors (Sox9) towards endocrine progenitors (Neurog3) and committed endocrine cells (Neurod1, Chga, Chgb). Although scRNA-seq analysis of the in vitro cells failed to detect hormones, except for Sst (Fig. 4B, F and Fig. 6D), immunostaining identified insulin and glucagon-expressing cells (Fig. 3D, Fig. S5 and Fig. S6C). Failure to detect mRNA encoding hormones in the PP+MP aggregates might have been due to their lower expression levels or to the higher expression of three genes: Rbpj, a central regulator of Notch signalling, GSk3b, a negative regulator of the Wnt pathway, and Insm1, a suppressor of Neurod1 and insulin-secreting cells (Fig. 6C-E). Almost all the cells that were positive for Ins1, Gcg or Sst in the embryonic endocrine population at different stages of pancreas development were negative for Rbpj, but some were positive for Gsk3b. Moreover, the embryonic endocrine population contained very few cells positive for Rbpj (Fig. 6E). In contrast, most of the endocrine population in the PP+MP aggregates was positive for Rbpj and Gsk3b, suggesting that Rbpj impaired full endocrine differentiation (Fig. 6E).

To test whether the PP+MP aggregates grown in suspension have fewer alpha and beta cells in comparison to the embryo, the relation between the expression of Yap1, Chga and Gsk3b was first assessed (Fig. 7A). The absence of an outer support layer of the aggregate was thought to possibly affect mechanical signalling, hence perturbing the expression of the mechanoresponsive transcription factor Yap1. A recent work by Mamidi et al. ^38^, suggested that extracellular cues inactivate Yap1, which triggers endocrinogenesis. In all samples, the Yap1 expression profile in the pancreatic aggregates is similar to that found in the embryonic pancreas, with a negative relationship between Yap1 and Chga expression (Fig. 7A and Fig. S7C). However, endocrine cells from the PP+MP aggregates expressed higher levels of Gsk3b in comparison to those from the embryonic pancreas (Fig. 7A). The higher expression level of Gsk3b might have prevented endocrine cells from further differentiating into insulin- and glucagon-secreting cells.

**Figure 7.**
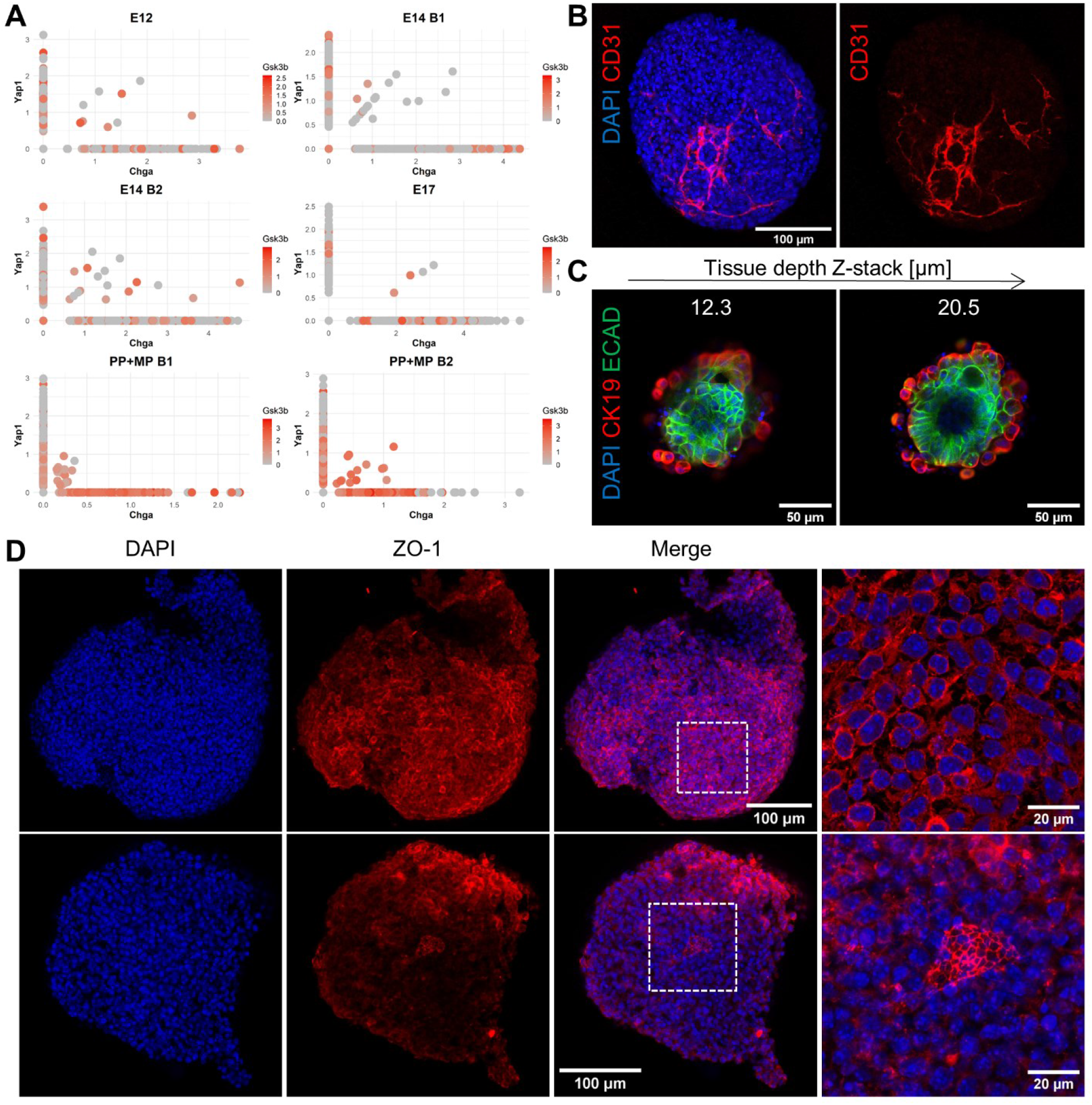
PP+MP aggregates do not lack mechanical cues for generating hormone-producing cells. **A.** Gene expression of Chga, Yap1 and Gsk3b in each sample. Each dot represents a cell, x and y axes show the normalized expression of Chga and Yap1, respectively. The normalized expression level of Gsk3b is indicated by the colour intensity. **B-C.** Confocal images of PP+MP aggregates embedded in 6.6% gelatin crosslinked with 5% microbial transglutaminase (mTG). **B.** The embedded aggregates were immunofluorescently stained for CD31, which marks the vessel-like network seen in the aggregate. **C.** The embedded aggregates were immunofluorescently stained for ECAD and CK19 (right), showing the epithelial and ductal regions at different tissue depths (Z-stacks). **D.** The non-gelatin-embedded PP+MP aggregates were immunofluorescently stained for zonula occludens-1 (ZO-1), which shows the tight junctions in the aggregates. The right-hand column presents higher-magnification images of the dashed square region in the merged image. Nuclear staining with DAPI is in blue.

To further investigate whether growing pancreatic aggregates in suspension can explain the disparity between in vitro and in vivo endocrine differentiation, PP+MP aggregates were embedded in 6.7% gelatin crosslinked with 5% microbial transglutaminase (mTg, Fig. 7B-C and Methods), an FDA-approved enzyme that catalyses formation of covalent bonds between lysine and glutamine groups of proteins and gelatin polymers ^64–66^. The gelatin-mTG hydrogel was used here to provide mechanical support to the aggregate, but also since it has been shown to improve cell maturation and long-term culture ^64,66,67^. The gelatin-embedded PP+MP aggregates contained a CD31-positive vessel-like network, as well as ductal regions marked by ECAD and CK19 (Fig. 7B-C and Video S1). However, insulin- and glucagon-expressing cells were not detected, suggesting that mechanosignalling was not a factor limiting in vitro formation of hormone-producing cells.

To determine whether the non-gelatin-embedded PP+MP aggregates maintained an organized epithelial structure, they were immunofluorescently stained for zonula occludens-1 (ZO-1, Fig. 7D). ZO-1 staining provides insights into the apical-basal polarity of tissues by visualizing tight junctions, which are crucial for the establishment of barrier function and apical-basal polarity ^68^. ZO-1 staining revealed intact and organized tight junctions primarily localized at the apical membrane of epithelial cells within the PP+MP aggregates, indicating the establishment of apical-basal polarity (Figure 7D). However ring-like epithelial structures exhibiting lumen were not demonstrated, as opposed to human PSC-derived ductal pancreatic organoids ^17,18^.

### Notch signalling pathway inhibition can rescue endocrine differentiation to alpha and beta cells in PP+MP aggregates

The single-cell RNA-seq analysis presented in Figure 6, which compared the endocrine populations between the embryo datasets and the PP+MP aggregates, identified a difference in the expression of Gsk3b and Rbpj and raised the possibility that higher expression of Rbpj in the PP+MP aggregates impaired full endocrine differentiation. Therefore, the influence of inhibition of Notch and GSK3 or of removal of the Wnt activator R-spondin1 on the endocrine differentiation of PP+MP aggregates was assessed. The expression of progenitor, endocrine and exocrine genes of interest were compared using RT-qPCR either after altering the aggregation medium throughout the 3D culture period or after subjecting the aggregates to different aggregation media from day 4 AA onwards. The control was that PP+MP aggregates were subjected to organoid medium without any alteration during the 3D culture (Fig. 8).

**Figure 8.**
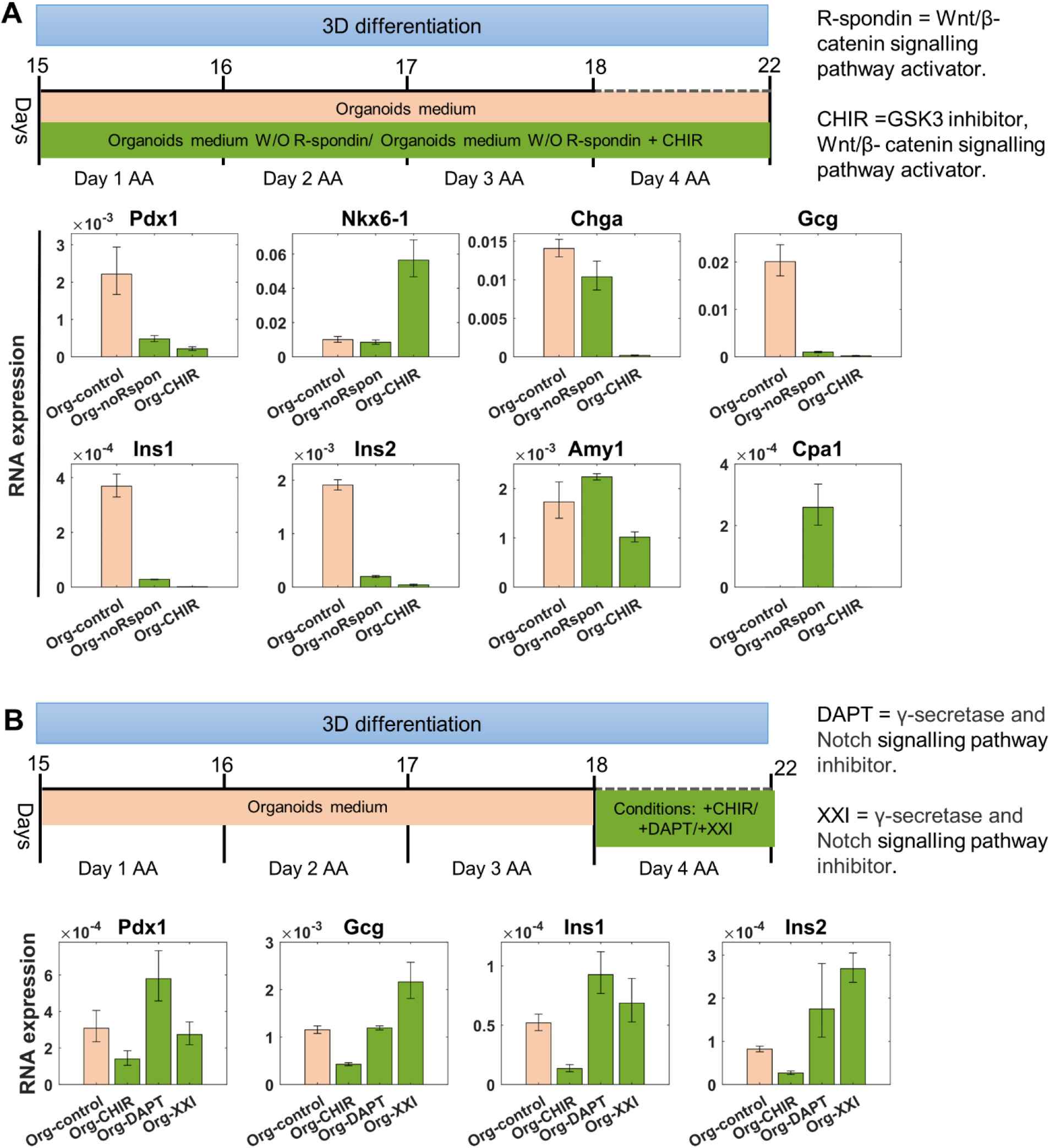
Inhibition of the Notch signalling pathway increased the expression of hormone-producing cells. **A-B.** A schematic illustration of PP+MP preparation with organoid medium alteration and RT-qPCR assessment of relative RNA expression of select pancreatic marker genes. Data are presented as averaged (n=96) aggregates per condition. Error bars indicate the SD. **A.** On day 14, 1000 mixed PPs and MPs cells were plated in each well of a U-bottom 96-well plate in organoid medium depleted from mouse R-spondin1 (Wnt activator) or in organoid medium depleted from mouse R-spondin1 and supplemented with 10 µM Chiron (GSK3 inhibitor). **B.** During the first 3 days AA, PP+MP aggregates were grown in organoid medium. From day 4 AA until the end of the differentiation, the aggregates were grown in organoid medium supplemented with either 10 µM Chiron, 1 µM DAPT (γ-secretase and Notch signalling pathway inhibitor) or 1 µM XXI (Compound E, γ-secretase and Notch signalling pathway inhibitor).

Removal of R-spondin1 from the aggregation medium, elevated the expression of exocrine markers amylase and Cpa1 (Fig. 8A) and lowered the expression of hormone genes in comparison to the control medium. Addition of CHIR, a GSK3 inhibitor, to the aggregation medium which was depleted of R-spondin1, lowered the expression of both endocrine and exocrine genes, excluding Nkx6-1 (Fig. 8A). Moreover, aggregates treated with CHIR seems to be larger than those grown under other conditions (data not shown), suggesting that addition of CHIR increased the progenitor pool.

Notch inhibition was tested with DAPT or XXI, which were added to the aggregation medium from day 4 AA. This modification of the aggregation medium was made 4 days AA, to allow the PP+MP aggregates to establish endocrine progenitors, as was indicated from the single cell RNA-seq analysis, which could then, with the right signalling environment, serve as fertile ground for full endocrine differentiation. Notch blocking by the two inhibitors increased the expression of Ins1 and Ins2, however only the addition of XXI to the medium led to an elevation in the expression of glucagon (Fig.8B). Addition of CHIR to the aggregation medium 4 days AA did not result in elevation in the hormone genes (Fig. 8B), suggesting, as the single-cell RNA-seq implied, that Rbpj, a Notch activator, impaired the full endocrine differentiation of the PP+MP aggregates, which was rescuable by Notch inhibition.

### Unique in vitro populations share similarities with the embryo

The proliferating cell cluster (cluster 2) was a distinct cluster among the in vitro aggregates, which scarcely contained embryonic cells (Fig. S8A). The identity of this cluster was not clear, therefore the gene profile proximity between cluster 2 and other known cell types was calculated. The 12 clusters found for the in vitro aggregate datasets (Fig. 4A) were projected on the aggregate-embryo integrated UMAP plot (Fig. S8B). The proliferating cell cluster mainly contained cells from four in vitro clusters (Fig. S8A), namely PO0, PO2, PO4, and PO5, which accounted for 99% of the proliferating cell cluster and was surrounded by known embryonic clusters (Fig. S8B). To assess the similarity between the four in vitro clusters in the proliferating cluster and clusters that had embryonic cells (14 clusters, Fig. S8), mutual information (MI) was calculated between genes and clusters. The detected informative genes (Fig. S8B and Table S3) were used to calculate the cosine similarity between the 14 known cell types and the four in vitro clusters (Fig. S8C and methods). The in vitro clusters displayed a significant similarity to erythroblast cells, with cosine similarity scores above 0.5 (similarity range was between −1 to 1, whereas −1 indicates that the conditions are opposite and 1 indicates maximal similarity), and to myeloid and immune cells, with approximately 0.45 cosine similarity scores (Fig. S8C). Furthermore, PO0 primarily resembled neuronal cells, and also exhibited a mild similarity to pancreatic progenitors, endocrine progenitors, endocrine cells and ENS neurons and endocrine (Figure S8C). PO2 showed similarity to neuronal cells, and mild similarity to acinar cells and ENS neurons and endocrine cells. PO4 and PO5 displayed similarity to ENS neurons and endocrine cells, alongside mild similarity to acinar cells, while PO5 also exhibited a mild similarity to neuronal cells. Furthermore, the cosine similarity profile of the in vitro clusters corresponded to their positions in the integrated UMAP (Fig. S8B). The proliferating cell cluster was surrounded by myeloid, erythroblast, immune cells and acinar cells. Additionally, some cells from PO0 and PO2 were projected onto the progenitor and endocrine progenitor cluster positions of the integrated UMAP (Fig. 5D and Fig. S8B). All the above suggests that the in vitro clusters represent modified versions of these cell types in the in vitro environment.

## Discussion

In this study, a well-defined in vitro culture system of mESC-derived progenitors was designed to mimic development of the mouse embryonic pancreas. The first step relied on differentiation of mESCs to PPs using a protocol adapted from human ESC differentiation to PPs ^12^. The main changes in the differentiation protocol were the removal of Matrigel and advancement of mESCs to the epiblast state before differentiation. The initial pluripotency state of the cells (2i, SL and EpiSCs) prior differentiation was shown to determine the capacity of the cells to differentiate to definitive endoderm and to dictate the subsequent PP differentiation yield. As compared to mESCs grown in 2i and SL prior differentiation, EpiSCs exhibited higher Sox17 expression at the DE stage and had a PP differentiation yield of above 86%, as determined by the coexpression of Pdx1 and Nkx6-1 and other pancreatic genes. This yield was remarkable compared to previously reported yields of 70% or less, even after a cell sorting step ^12,35–38^. Furthermore it emphasized the responsiveness of EpiSCs to activin and their capacity to commit to DE, which is a major step in ensuring a successful PP differentiation protocol ^69^.

Two recent works made advances in achieving expandable PPs from human PSCs, either based on growing expandable human PPs on feeder cell layers ^70^ or as PP-spheroids in Matrigel ^19^. The long-term maintenance and expansion of pancreatic progenitors represents a significant breakthrough in biomedical research and regenerative medicine. It opens new avenues for obtaining an unlimited supply of pancreatic organoids and the differentiation of these progenitors to various pancreatic cell types, including beta cells. The mouse EpiSC-derived PPs described here survived a freeze-thaw cycle (data not shown), however further work is still needed to establish the optimal conditions for robust PP expansion. Furthermore, implementation of the recent advances in the human expandable PPs on the mouse system, might shed light on the differences between human and mouse pancreatic systems.

After establishing a reproducible protocol for PP production, efforts were made to mimic the 3D pancreatic microenvironment of the developing mouse pancreas. To this end, EpiSC-derived pancreatic and mesodermal progenitors were mixed in proportions similar to those measured in the embryo ^44^ to form an aggregate. The MPs supplied the PPs with mesodermal cues, but also cellular interactions via specialized cell types, such as endothelial and mesenchymal cells, which are essential for pancreas differentiation and maturation in vivo. It has been shown that growing progenitors on organ-matched mesenchyme elicits their expansion and self-renewal and, importantly, gives rise to glucose-sensing and insulin-secreting cells when transplanted in vivo ^71^. Here, aggregation of EpiSC-derived PPs with EpiSC-derived MPs without the use of Matrigel, resulted in aggregates containing a range of cell types, including pancreatic progenitors, epithelial, acinar, endocrine, mesenchymal and endothelial cells. Moreover, insulin and glucagon expression were detected in immunostaining analyses, indicating the presence of beta and alpha cells, respectively.

A single-cell transcriptomic comparison between PP+MP aggregates, aggregates consisting solely of PP cells and 3D culture of PPs mixed with Matrigel was then performed to gain a better understanding of the benefits provided by the mesodermal compartment within the pancreatic aggregates. The scRNA-seq extended the depth of the in vitro system analysis and revealed the diverse cell populations that existed in the 3D in vitro systems. Notably, the PP+MP aggregates displayed a distinct profile with a more abundant representation of endocrine cells compared to PP and PP+Matrigel aggregates. Moreover, PP+MP aggregates demonstrated higher endothelial cell percentage in comparison to the other in vitro samples. These finding suggests that endothelial cells are important for endocrine differentiation, as indicated by previous studies which showed that endothelial cells repress acinar differentiation whilst increasing endocrine differentiation ^48,72^.

The cell population comparison between the three in vitro systems further indicated that although PP+MP aggregates showed no deficiency in ECM proteins, the addition of Matrigel to PPs induced the expression of ductal markers. Recent studies showed that Matrigel triggered epithelial morphogenesis in vitro, as demonstrated, for example, by the emergence of a neural tube shape after embedding aggregates of mouse ESCs in Matrigel ^73^, epithelia dome-shaped structures of human PSC monolayers grown in the presence of Matrigel in ^74^ and epithelial ring organization with apical-basal cell polarity of human PSC-derived ductal pancreatic organoids cultured with Matrigel ^17,18^. In the present study, the hydrogel-free PP+MP aggregates expressed the tight junction marker ZO-1 but did not exhibit lumen- or ring-like structures similar to those observed in the ductal pancreatic organoids. In contrast, the gelatin-embedded PP+MP aggregates showed improvement in the epithelial organization, suggesting that gelatin, by imposing biophysical constraints, together with the mesodermal compartment, providing biochemical signals, can compensate for the Matrigel epithelial patterning regulation. Overall, the discrepancy in cell type composition between the in vitro systems highlights the unique characteristics and potential of PP+MP aggregates.

Some discrepancies were found between the immunofluorescence staining and the scRNA-seq results, mainly related to insulin and glucagon, which were not detected at the mRNA level. This may have been due to the pooling of single cells from hundreds of aggregates for the scRNA-seq analysis, which might have interfered with the detection of rare cell types and genes expressed at low levels, e.g., hormone-secreting cells.

The comparison of single-cell transcriptomes of the 3D in vitro samples to the embryonic data collected at E12, E14 and E17 demonstrated that the in vitro aggregates succeeded in recapitulating many of the cell populations residing in the pancreas. Remarkably the PP+MP aggregates had an endocrine compartment similar in its size to that of the E14 and E17 pancreas and endothelial compartment similar to that of the E14 pancreas. On the other hand, the PP and PP+Matrigel samples exhibited dominant neuronal populations. Resemblance to the embryonic pancreas was not only in the relative size of the endocrine population but also in the differentiation path from the progenitors to endocrine cells, particularly in the PP+MP sample. Yet the PP+MP aggregates did not entirely mirror the embryo and contained a population (cluster 2) that was distinct from the embryonic pancreatic cells. Although this population was mainly composed of in vitro cells that did not cluster with the embryonic pancreatic cells, they exhibited a gene profile that was close to known clusters as erythroblast, myeloid, progenitors and endocrine cells. This gene profile discrepancy between the pancreatic aggregate cells and the in vivo cells, might reflect differences in the main signalling pathways driving pancreas development, i.e., Notch, Wnt and Hippo pathways. A recent work by Mamidi et al. ^38^, suggested that extracellular cues inactivating Yap1 trigger endocrinogenesis. In the current in vitro system, although the PP+MP aggregates were grown in suspension and lacked the mechanical support typically provided by Matrigel, most of the cells that were positive for Chga were also negative for Yap1, indicating that the in vitro progenitors can produce endocrine cells. The endocrine cells of the PP+MP aggregates also expressed higher level of Gsk3b, a Wnt repressor, and of Rbpj, a central regulator of Notch signalling, in comparison to the embryo. Notch signalling helps maintain progenitor proliferation and prevents premature differentiation to ductal and endocrine fate ^58,75^. Wnt signalling regulates pancreatic specification and patterning during different stages of pancreas development ^58,59^. On the one hand, Wnt inhibition is necessary for endocrine differentiation ^76^, but, on the other hand, deletion of Wnt-related genes has been found to reduce the proportion of pancreatic progenitors, which subsequently reduces the number of both endocrine and exocrine cells ^58^. In light of an earlier report that Gsk3b overactivation results in decreased pancreatic beta cell proliferation and mass ^77^ and that high levels of Notch signalling lead to the repression of Neurog3, preventing endocrine cell fate determination ^58^, the ability of Gsk3b and mainly Rbpj to limit endocrine cells from further differentiating into insulin- and glucagon-secreting cells was evaluated here. Exposure of PP+MP aggregates to Notch inhibitors increased the expression of insulin and glucagon, however addition of GSK3 inhibitor to the aggregate growth medium did not increase the expression of hormone genes, indicating that indeed Rbpj impaired full endocrine differentiation in the PP+MP aggregates. Overall, the transcriptomic analyses revealed the similarities and differences between the pancreatic aggregates and the mouse embryonic pancreas and highlighted signalling pathways and mechanisms that may govern pancreas progenitor differentiation and development.

To further determine whether the lack of mechanical cues generally provided by Matrigel, prevented the emergence of insulin- and glucagon-producing cells in the in vitro system, the PP+MP aggregates were embedded in a gelatin-mTG hydrogel, which constrains cells in space. However, cells positive for insulin or glucagon were not detected in the gelatin-embedded aggregates, implying that mechanical cues are either already present via the MP counterpart of the aggregate or that they are not the key determinant of full endocrine differentiation in this pancreatic in vitro model. Nevertheless, the gelatin-embedded aggregates were highly vascularized and displayed epithelial ductal regions, which are important for in vitro maturation and indicate correct development. Achieving such vascularization is an advancement in tissue engineering, since vascularization is important for growth, prevents necrosis as the aggregates increase in size and prevents premature differentiation ^78^.

Until recently, the study of embryo and organ development has been restricted to animal embryos and fetal tissues. Although organogenesis can be observed in the embryo, these animal models and material sources are limited, and their use presents ethical concerns. The increasing utilization of human and mouse stem cell-derived organoids has significantly expanded the study of mammalian development and promoted regenerative and therapeutic medicine, whilst reducing the use of animals. Nevertheless, most organoids, and specifically pancreatic organoids, rely on the use of Matrigel, which limits reproducibility and full definition of an in vitro 3D system. The present study aimed to maximize in vitro resources to construct a new in vitro system by mixing endoderm- and mesoderm-derived progenitors in a defined proportion. In the future, these mixtures can be explored in the in vitro system, and might shed light on the important interactions between the endoderm- and mesoderm-derived progenitors. The novel system closely recapitulated key elements of mouse embryonic pancreas and bears the potential to serve as a tuneable model for the study of mammalian pancreas development and pancreas diseases.

## Methods

### Cell culture

ES-E14TG2a (ATCC CRL-1821) mouse embryonic stem cells were grown in sterile flasks pre-coated with 0.1% porcine skin Type A gelatin (Sigma-Aldrich, G1890) in pluripotency medium referred to as serum and LIF (SL), comprised of GMEM (Gibco) supplemented with 1% non-essential amino acids (100X Gibco), 1 mM sodium pyruvate (Biological Industries), 1% GlutaMAX (100X Gibco), 0.1 mM β-mercaptoethanol (Gibco), 10% foetal bovine serum (FBS) (HyClone, Thermo Fisher Scientific), and 10 ng/ml LIF (R&D Systems). Cells were maintained at 37 °C in 5% CO_2_. Medium was changed daily, and cells were passaged with trypsin (Biological Industries) every 2-3 days, as necessary, up to a total of 30 passages.

### 2i

2i was comprised of basal differentiation medium NDiff®227 (Takara), hereby referred to as N2B27, supplemented with 1 μM MAPK inhibitor PD0325901 (Sigma-Aldrich, referred as PDO3) and 3 μM GSK3β inhibitor CHIR99021 (Sigma-Aldrich, referred to as Chiron). Cells grown in 2i medium are in a stringent pluripotency environment and are considered to be at ground-state pluripotency ^32^.

### Epiblast stem cell and Epi-medium

ES-E14TG2a were grown in plastic tissue-culture flasks coated with 5 μg/ml plasma fibronectin (F0895, 1 mg/ml, Sigma-Aldrich) in Dulbecco′s phosphate-buffered saline (DPBS with calcium and magnesium, Sigma-Aldrich, D8662). mESCs were grown in Epi-medium which was comprised of N2B27 supplemented with 12 ng/ml FGF2 (Peprotech) and 25 ng/ml Activin A (Peprotech), for at least four passages, to generate EpiSCs ^42^.

### Differentiation of mESCs to pancreatic progenitors (PPs)

The differentiation protocol below is a variation of one previously used to derive PPs from human pluripotent stem cells ^12^. Prior to differentiation, mESCs were grown in either SL or 2i or as EpiSCs in Epi-medium. Cells grown in SL and 2i were grown in sterile flasks pre-coated with 0.1% gelatin, whereas flasks with EpiSCs were pre-coated with 5 μg/ml fibronectin.

Cells were plated at a density of 5×10^4^ cells/cm^2^ in their respective growth medium (SL, 2i or Epi-medium) 24 h prior to differentiation (day 0). The backbone medium for the first 5 days of differentiation was BE1: MCDB131 (Gibco) supplemented with 0.8 g/L cell-culture-tested glucose (Sigma-Aldrich), 1.174 g/L sodium bicarbonate (Biological Industries), 0.1% fatty acid-free (FAF) bovine serum albumin (BSA) (Sigma-Aldrich) and 2 mM GlutaMAX. The backbone medium for the subsequent 9 days of differentiation was BE3: MCDB131 supplemented with 0.44 g/L glucose, 1.754 g/L sodium bicarbonate, 2% FAF-BSA, 2mM GlutaMAX, 4.4 mg/L-ascorbic acid (Sigma-Aldrich) and 0.5x insulin-transferrin-selenium-ethanolamine (Gibco). Cells undergoing differentiation were cultured at 37 °C in a 5% CO_2_ incubator and medium was changed daily. On day 1, cells were washed with DPBS and incubated with BE1 supplemented with 3 µM Chiron (CHIR99021, Sigma-Aldrich) and 100 ng/mL activin A. On day 2, medium was replaced by BE1 with 100 ng/mL activin A for 2 days. On day 4, medium was changed to BE1 with 50 ng/mL keratinocyte growth factor (KGF also known as FGF7, Peprotech) for 2 days. On days 6-9, cells were cultured in BE3 medium containing 0.25 µM SANT-1 (Sigma-Aldrich), 2 µM retinoic acid (Sigma-Aldrich), 200 nM LDN-193189 (Sigma-Aldrich) and 500 nM PDO3 (PD0325901 Sigma-Aldrich). On days 10–14, the cells were cultured in BE3 supplemented with 50 ng/mL FGF10 (R&D systems), 330 nM Indolactam V (Cayman), 10 µM SB431542 (Tocris) and an additional 16 mM glucose. An outline of the differentiation protocol is presented in Fig. 1A.

### Mesoderm progenitors (MPs)

The protocol for differentiation of EpiSCs to mouse embryo caudle epiblast (CE)-like cells is detailed in ^41,42^. Briefly, EpiSCs were differentiated to Epi-CE, which resemble the caudle epiblast of the mouse embryo and contain the different mesoderm compartments: lateral plate mesoderm, intermediate and paraxial mesoderm, hence we refer to these cells here as mesoderm progenitors. EpiSCs were plated at a density of 5 × 10^4^ cells/cm^2^ in a flask pre-coated with 5 μg/ml fibronectin and grown in Epi-medium for the first day (day 0). On Day 1, medium was replaced with N2B27 supplemented with 20 ng/ml FGF2 and without activin A. On day 2, N2B27 was supplemented with 3 μM Chiron and 20 ng/ml FGF2.

### Aggregate preparation

#### PP+MP aggregates

PPs and MPs in adherent culture were dissociated from culture flasks with accutase (STEMCELL Technologies) on days 14 and 3 of differentiation, respectively. Cells (n=1000) at a ratio of 58% PPs and 42% MPs were plated in each well of a U-bottom 96-well plate in 50 μL organoid medium comprised of Dulbecco’s Modified Eagle Medium/Nutrient Mixture F-12 (DMEM/F-12(HAM)1:1, Biological Industries) with 10% knockout serum replacement (KSR, Gibco), 1% penicillin-streptomycin (PS, Biological Industries), 0.1 mM β-mercaptoethanol (Gibco), 16 nM phorbol myristate acetate (PMA, Sigma-Aldrich), 5 μM ROCK inhibitor (Thazovivin, Sigma-Aldrich), 25 ng/mL epidermal growth factor (EGF, Sigma-Aldrich), 500 ng/ml mouse R-spondin1 (R&D systems), 2.5 U/mL heparin sodium salt (Sigma-Aldrich), 25 μg/mL acidic fibroblast growth factor (aFGF, R&D systems) and 100 ng/mL FGF-10 ^23^. After 48 h, 100 μL of the same medium was added to each well for an additional 48 h. On day 4 AA, the aggregates were transferred to a 6-well plate and placed on an orbital shaker operating at 110 RPM (RCF of 0.11808g and shaking diameter of 10mm) for an additional 4 days (total 22 days of differentiation, Fig. 3B). Wells were supplemented with 3 ml organoid medium, of which 1 ml was changed every 2 days from day 4 AA until the end of differentiation.

Aggregate cell ratio justification: At embryonic stages E12.5 and E14.5, the pancreas was composed of 58% epithelial cells, 37% mesenchymal cells and 5% endothelial cells (Fig. 5C in ^44^). The mesoderm gives rise to mesenchyme, endothelium and blood cells ^4,79^, hence MPs represent the mesenchymal and endothelial cells and PPs represent the epithelial compartment of the pancreas embryo, requiring in a ratio of 58% PPs and 42% MPs in an aggregate (Fig. 3A).

#### PP aggregates

PPs in adherent culture were dissociated from culture flasks with accutase on day 14 of differentiation. 1000 cells were plated in in each well of a U-bottom 96-well plate in 50 μL organoid medium. Culture and medium change were the same as for PP+MP aggregates.

#### PP+Matrigel

Generation of PP aggregates in Matrigel was based on a protocol previously described for human PP-spheroids ^19^. PPs in adherent culture were dissociated from culture flasks with accutase on day 14 of differentiation. One part of 4×10^6^ cells/ml was mixed with 3 parts of Matrigel (Matrigel® Growth Factor Reduced (GFR) Basement Membrane Matrix, Phenol Red-free, LDEV-free, Corning® 356231). An example for cells-Matrigel mixture volumes is 40 µl of 4×10^6^ cells/ml mixed with 120 µl Matrigel. Drops of 40 µl cells-Matrigel mixture (40,000 cells per drop) were slowly dispensed onto the center of each well of pre-warmed 4-well plates (Nunc cell culture, Thermo Fisher Scientific), to form a 3D dome. The plates were incubated at 37 °C for 10 min to allow the Matrigel to solidify and then 500 µl organoid medium was added to each well. Organoid medium was changed every 3 days.

### RNA extraction and real-time quantitative polymerase chain reaction (RT-qPCR)

RNA was extracted at different time points during EpiSC differentiation to PPs using the RNeasy extraction kit (Qiagen), according to the manufacturer’s instructions. RNA concentration was measured with a NanoDrop (NanoDrop One, Thermo Fisher Scientific). cDNA was synthetized using the high-capacity cDNA Reverse Transcription Kit (Applied Biosystems, Thermo Fisher Scientific), according to the manufacturer’s instructions. RT-qPCR was performed using Fast SYBR Green Master Mix (Applied Biosystems, Thermo Fisher Scientific), according to the manufacturer’s instructions. The gene-specific primers that were used are listed in Table 1. The reaction was performed in a QuantStudio1 (Applied Biosystems, Thermo Fisher Scientific), in technical triplicates. All experiments were performed in biological triplicates. Expression values were normalized against the housekeeping gene Ppia (2^−Δct^). The results are presented as the average across biological replicates (2^−mean(Δct)^) with the standard error of the mean (SEM, 2^−(mean(Δct)±sem(Δct))^).

**Table 1.**
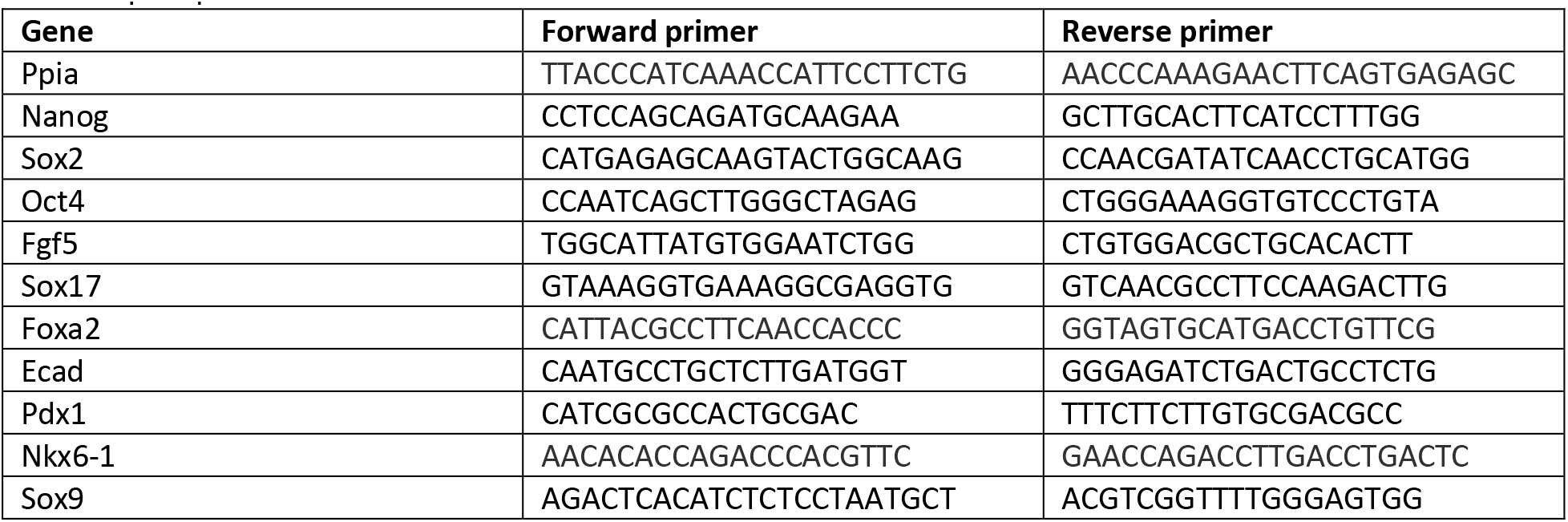
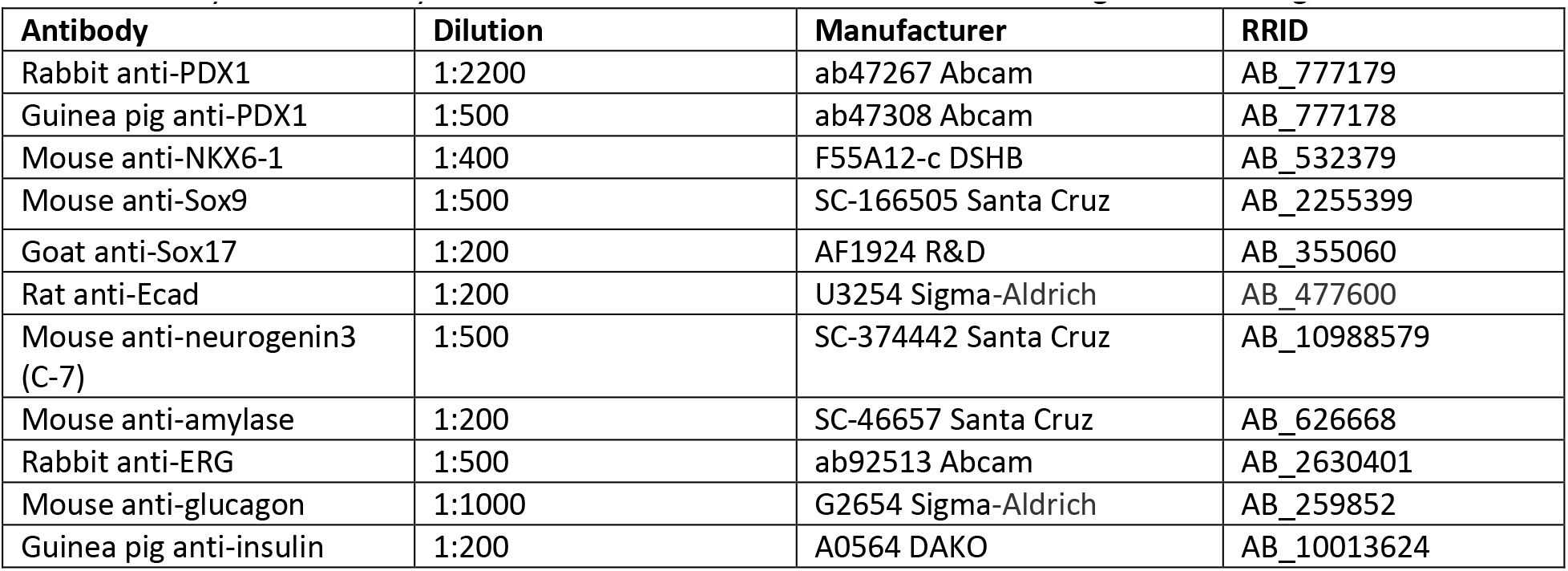
qPCR primer list.

### Cell viability assay

The 3-(4,5-dimethylthiazol-2-yl)-2,5-diphenyltetrazolium (MTT) assay based on the conversion of MTT to blue formazan crystals by viable cells, was used to measure cell viability. Briefly, 24 h after mESCs were grown in either 2i, SL or as EpiSCs in Epi-medium (day 0) and on days 4, 6 and 10 during PP differentiation, the medium was removed and replaced with a fresh MCDB131 medium with 0.5 mg/ml MTT (Glentham Life Sciences, GC4568 Corsham, UK). After incubation at 37 °C for 2 h, the medium was discarded and 100 µl DMSO was added to each well. The plate was incubated at room temperature for an additional 2 h and absorption was determined in a microplate reader (Synergy H1, BioTek) at λ = 570 nm/650 nm after automatic subtraction of background readings. Cell viability was expressed as the percentage of total EpiSCs at the relevant time point during PP differentiation.

### Immunofluorescence and confocal imaging

Cells and aggregates were washed with phosphate-buffered saline (PBS) 1X (Gibco) and fixed in 4% paraformaldehyde (PFA) (ChemCruz, Santa Cruz Biotechnology) for 15 min at room temperature (RT). Samples were washed and permeabilized for 15 min with PBST:PBS 1X + 1mM CaCl_2_, supplemented with 0.5% bovine serum albumin (BSA) (Millipore) and 0.5% Triton X-100 (Bio-Lab Ltd.) before overnight incubation at 4 °C with primary antibodies diluted in PBST. Antibody used are listed in Table 2. The following day, samples were washed with PBST and incubated at room temperature for 2 h with secondary antibodies and 4′,6-diamidino-2-phenylindole (DAPI) (Sigma-Aldrich) diluted in PBST. Samples were washed with PBS 1X and stored in fresh PBS 1X at 4 °C until imaging. Aggregates were mounted on microscope slides by pipetting them in 15 μl droplets of Fluoromount-G (SouthernBiotech, 0100-01) and then covered with a microscope cover glass (No. 1.5H 24 mm x 60 mm) with an adhesive spacer (Silicone isolators, press-to-seal, Sigma-Aldrich). Cells and aggregates were imaged using a confocal microscope (LSM700, Ziess) and Fiji software (ImageJ) was used to process and analyse the images.

**Table 2.**
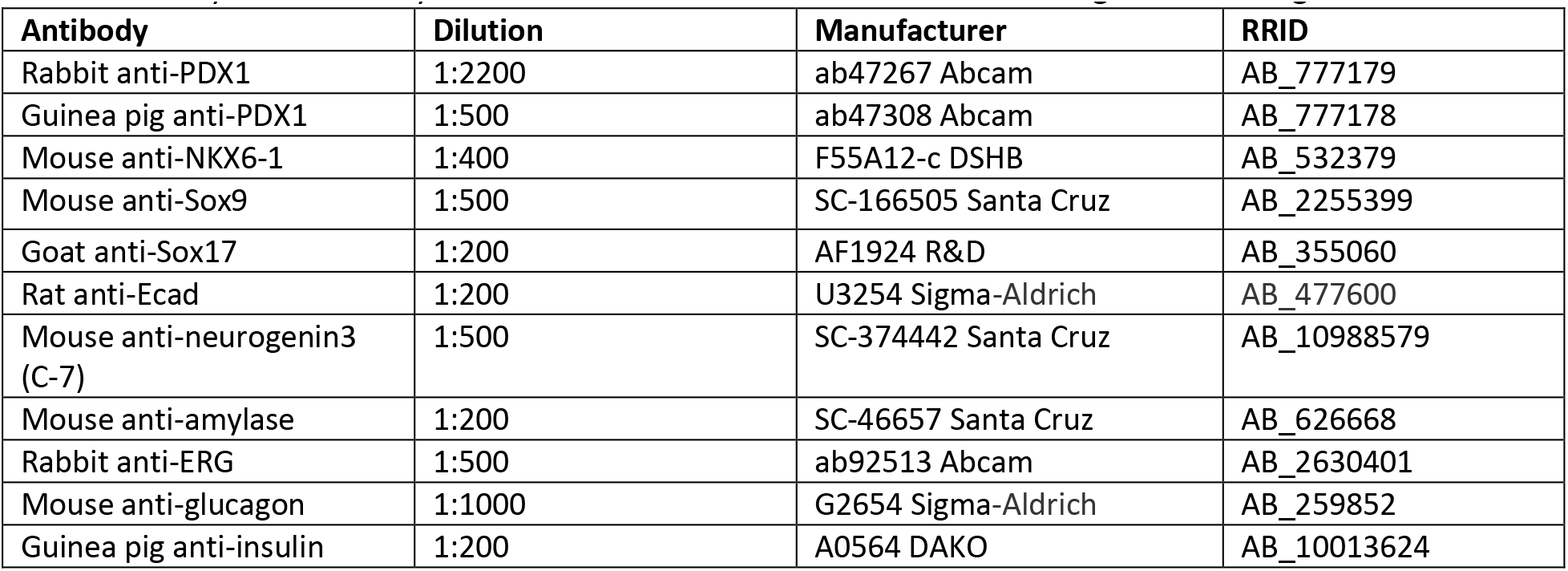
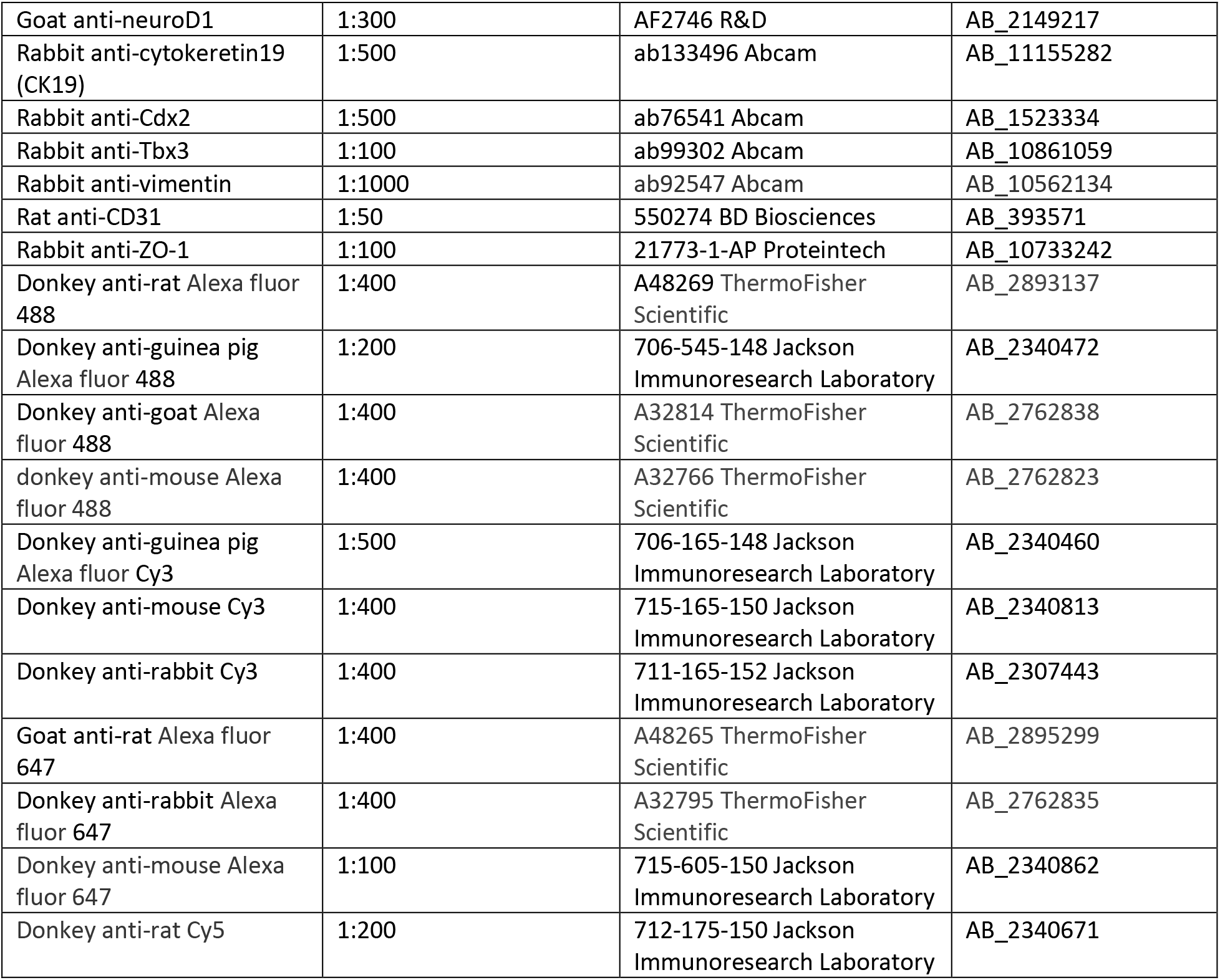
Primary and secondary antibodies used for immunofluorescence staining of cells and aggregates.

### Flow cytometry

To assess the differentiation of EpiSCs to PPS, flow cytometry was performed on day 14 using mouse anti-NKX6-1 and rabbit anti-PDX1 antibodies (Table 2). EpiSC-derived PPs were fixed in 4% PFA for 15 min. Cells were centrifuged at RT for 3 min at 300 *g* to remove the fixation solution, and then washed in PBST. The cells were incubated with anti-PDX1 and anti-NKX6-1 primary antibodies overnight at 4 °C. The following day, the cells were centrifuged at RT for 5 min at 300g, washed in PBST and centrifuged at RT for 5 min at 300g. Next, the cells were incubated with donkey anti-mouse Alexa fluor 647 (1:100, Jackson Immunoresearch Laboratory) and donkey anti-rabbit Cy3 (1:400, Jackson Immunoresearch Laboratory) secondary antibodies for 2 h at RT, in the dark. Cells were washed in PBST, centrifuged at RT for 5 min at 300g and resuspended in PBS 1X. Control samples were incubated with secondary antibodies only. Cells were analysed using the BD FACS Aria-IIIu cell sorter (BD) and the data were analysed with FlowJo analysis software (BD).

### Single-cell RNA sequencing (scRNA-seq)

On day 22 (day 8 AA), PP+MP aggregates (2 biological replicates), PP aggregates and PP+Matrigel were dissociated into single cells. PP+MP and PP aggregates were dissociated with accutase, whereas PP+Matrigel were dissociated with TrypLE Select 1X (Gibco). A 10x Genomics single-cell transcriptomic service was used to sequence the cells of the aggregates. For batch 1, a single-cell RNA sequencing library was generated for 3,121 cells from PP+MP aggregates, while for batch 2, three libraries were generated for 6,602 cells of PP+MP aggregates, 7,575 cells of the PP aggregate and 4,240 cells of the PP+Matrigel sample. Briefly, after applying various filtering steps, a total of 1456 and 3602 cells of PP+MP aggregates from batch 1 and batch 2 respectively, 5,056 cells of PP aggregate and 3,485 cells of PP+Matrigel were retained for subsequent analysis. To group cells by cell types, the graph-based clustering R package Seurat v4.3.0 ^80^ and the batch correction R package Canek version 0.2.1 ^81^ was used, followed by visualization using Uniform Manifold Approximation and Projection (UMAP) dimensionality reduction technique.

#### Library preparation and data generation

One RNA single-cell library for batch 1 and 3 RNA single cell libraries for batch 2 were prepared at the Technion Genomics Centre, according to the 10x manufacturer’s protocol (Chromium Next GEM Single Cell 3’ Library & Gel Bead Kit v3.1, PN-1000121), using 7,000 and 15,000 input cells per sample for batch 1 and batch 2, respectively. Single-cell separation was performed using the Chromium Next GEM Chip G Single Cell Kit (PN-1000120). The RNA-seq data was generated on Illumina NextSeq2000, P2 100 cycles (Read1-28; Read2-90; Index1-10; Index2-10) (Illumina, cat no. 20046811). Cell Ranger version 6 and version 7.1 (for batch 1 and batch 2 respectively, 10x Genomics) was used to process raw sequencing data and the Seurat R package version 4.3.0 ^80^ was used to build the expression matrix. Gene expression was quantified by unique molecular identifiers (UMI) counts.

#### Mouse embryo single-cell RNA-seq data

The single-cell transcriptomic data of pancreatic aggregates was compared to transcriptomic data of the mouse pancreas collected on embryonic days 12, 14, and 17 ^8^. As detailed by Byrnes et al. ^8^, single-cell RNA-sequencing libraries were generated for 4,631 cells on E12, 9,028 cells on E14 (comprised of two independent batches E14_B1 and E14_B2), and 4,635 cells on E17. Cell and gene filtering, normalization and data integration between the in vitro and the embryo samples were all performed with R package Seurat version 4.3.0 ^80^.

#### Single-cell data clean up and quality control

Using Seurat R package version 4.3.0 ^80^, the expression matrix was cleaned as follows: 1) UMI counts: only cells that had 500-70,000 UMI counts were retained for further analysis, ensuring a sufficient sequencing depth for each cell. 2) Detected genes: only cells that had 500-8000 unique genes in a cell were retained for further analysis, to ensure that the reads were distributed across the transcriptome. 3) Mitochondrial gene expression: an upper limit of 20% mitochondrial genes (mitochondrial gene counts in a cell versus the total detected genes in a cell) in a cell was set, to reduce the likelihood that cells for further analysis were dead or stressed. 4) Gene filtering: a threshold of at least 5 cells containing more than 1 UMI of a gene was set. The number of cells and total genes after the clean-up are presented in Table 3. The aggregate data was processed with Seurat version 4.3.0 and batch correction was performed with Canek R package version 0.2.1 ^81^. First, the expression matrix was log-normalized and then scaled (linear transformed), such that the mean and the variance expression of each gene across cells was 0 and 1, respectively. Next, principal component analysis (PCA) was performed, from which the first 30 principal components (PCs) were selected for uniform manifold approximation and projection (UMAP) analysis and cell clustering, which is a graph-based clustering approach. Integration of the in vivo and in vitro samples was performed using the Seurat anchoring method ^80^. The integrated expression matrix was log-normalized and scaled. Next, PCA was performed and the first 30 PCs were selected for UMAP analysis and cluster finding.

**Table 3.** The number of cells in each sample and the total number of detected genes after data clean up.

| Sample | Cell number | Gene Number |
| --- | --- | --- |
| E12 | 4,412 | 20,293 |
| E14_B1 | 3,495 | 20,293 |
| E14_B2 | 4,309 | 20,293 |
| E17 | 2,241 | 20,293 |
| PP_org | 5056 | 20,293 |
| PP+Matrigel | 3485 | 20,293 |
| PP+MP_B1 | 1456 | 20,293 |
| PP+MP_B2 | 3602 | 20,293 |

Marker genes for all clusters was detected as genes with at least Log2(fold change) ≥1 and adjusted P-value ≤0.05 (Table S2). The GSEA MSigDB (https://www.gsea-msigdb.org/gsea/msigdb/) and Descartes (https://descartes.brotmanbaty.org/bbi/human-gene-expression-during-development/dataset/pancreas) databases were used to help assign identities for the 20 clusters according to their marker genes.

### Inhibition of Wnt and Notch during aggregate preparation

PP+MP aggregates were grown as detailed above with the following modifications to the organoid medium:

1. After aggregation, the aggregates were grown in organoid medium without mouse R-spondin1 (Wnt activator) or in organoid medium without mouse R-spondin1 but supplemented with 10 µM Chiron (GSK3 inhibitor).
2. During the first 3 days AA, aggregates were grown in organoid medium. From day 4 AA until the end of the differentiation, the aggregates were grown in organoid medium supplemented with either 10 µM Chiron, 1 µM DAPT (γ-secretase and Notch signalling pathway inhibitor, Tocris 2634) or 1 µM XXI (Compound E, γ-secretase and Notch signalling pathway inhibitor, AdipoGen Life Sciences AG-CR1-0081).

A total of 96 aggregates per condition were combined and averaged for RT-qPCR analysis.

### Mutual information between genes and clusters

To characterize cluster 2, which was named ‘proliferating cells’, the similarity between the in vitro cells in cluster 2 and all the clusters that contained embryonic cells (14 clusters), was assessed. Downstream analysis was implemented with the entire set of qualified genes (20,293) rather than with the genes restricted to the integration process (2,000 anchor genes). This step was performed as previously detailed ^41,82^. A mutual information (MI) technique ^83^ was used to select the informative genes related to the embryonic clusters and the major in vitro clusters assembling the ‘proliferating cells’ cluster (4 clusters). The MI between the clusters (denoted as Y) and genes (denoted as X) was computed as follows:

1. Calculation of cluster entropy:

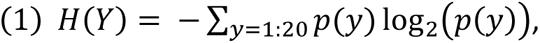 where *p*(*y*) is the probability of each cluster *y* = 1,2,…20.
2. Discretization of the gene expression values into ten bins and calculation of the conditional entropy *H*(*Y|X*) as follows:

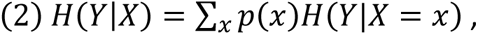 where *p*(*x*) is the probability of the discretized gene expression across the cell population and *H*(*Y|X* = *x*) is the cluster entropy given a specific gene expression value.
3. Computation of the MI between the clusters and each gene according to the equation below:

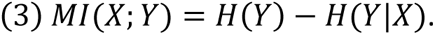
4. Setting of a threshold of the MI of all the genes and selection of the informative genes above this value (Fig. S8B).

Fig. S8B shows the MI between the genes and the clusters versus the number of genes. MI was set at 0.23, since at this point, the MI practically did not fluctuate with increments in gene number. This threshold, in which genes have MI above or equal to 0.23 (273 genes in total, Table S3), determines which genes are selected as input features for calculating the cosine similarity.

### Cosine similarity

Cosine similarity was used as a measure of similarity between gene expression of the informative 273 genes selected by MI (Table S3) in one cluster versus another cluster. The cosine similarity was calculated as follows:

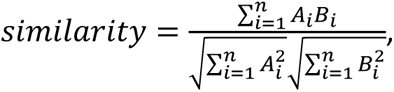

where A and B represent two clusters and A_i_ and B_i_ are the gene expression values in each condition (n=273). Similarity of −1 indicates that the conditions are opposite and 1 indicates maximal similarity.

### Aggregate embedding in gelatin

Gelatin hydrogel was covalently crosslinked with microbial transglutaminase (mTG) (MOO gloo TI transglutaminase, SKU:1203-50, modernist pantry) in the presence of calcium ions ^64^. Porcine skin gelatin (300 g Bloom type A, Sigma-Aldrich, G1890) was dissolved in PBS 1X at 50 °C and immediately passed through a 0.22 µm filter. mTG was dissolved in 10 mM HEPES buffer (Biological Industries) supplemented with 100 mM CaCl_2_ and sterilized by passing it through a 0.22µm filter.

On day 6 AA, PP+MP aggregates were embedded in 6.7% gelatin and 5% mTG, prewarmed to 37 °C. Plugs (50 µl) of gelatin-mTG with aggregates were pipetted into wells of a 24-well plate. To solidify the gelatin, the plate was incubated for 10 min at RT, followed by 30 min at 37 °C before the organoid medium was added.

## Acknowledgments

We would like to thank Janette Zavin for assistance with staining, cell culture and image illustrations, and Rita Shuhmaher and Dina Safina for assistance with cell culture and fruitful discussion throughout the project. We would also like to express our sincere appreciation to Galia Ben David for her invaluable assistance throughout the project. We are grateful to Orit Bar-Am and Hagit Sason for their support in conducting the MTT assay, and Liat Linde and her team from the Technion Genomics Center for performing the 3D in vitro pancreatic aggregate single-cell RNA-sequencing. Images in Figures S1A were created with BioRender.com.

## Author Contributions

Conceptualization: S.E. and S.L.; Methodology: S.E, N.S. and S.L.; Software: S.E.; Validation: S.E., V.R., O.G. and A.N.F.; Investigation: S.E., V.R. and O.G.; Resources: S.L.; Writing - original draft: S.E. and S.L.; Writing - review & editing: S.E., A.N.F., N.S., C.E.P. and S.L.; Supervision: S.L.; Funding acquisition: S.L. and C.E.P.

## Data and materials availability

All data needed to evaluate the conclusions in the paper are present in the paper and/or in the Supplementary Materials. The raw data for single-cell RNA sequencing of PP+MP aggregates (2 biological replicates), PP aggregates, and PP+Matrigel samples is accessible for download through the provided link: https://data.mendeley.com/datasets/6hj6t5jgnc/draft?a=b2bcb3cb-16c3-45b7-b603-7270502e8f05

