## Supplementary figures and images for "3D model of mouse embryonic pancreas and endocrine compartment using stem cell-derived mesoderm and pancreatic progenitors"

### Video S1

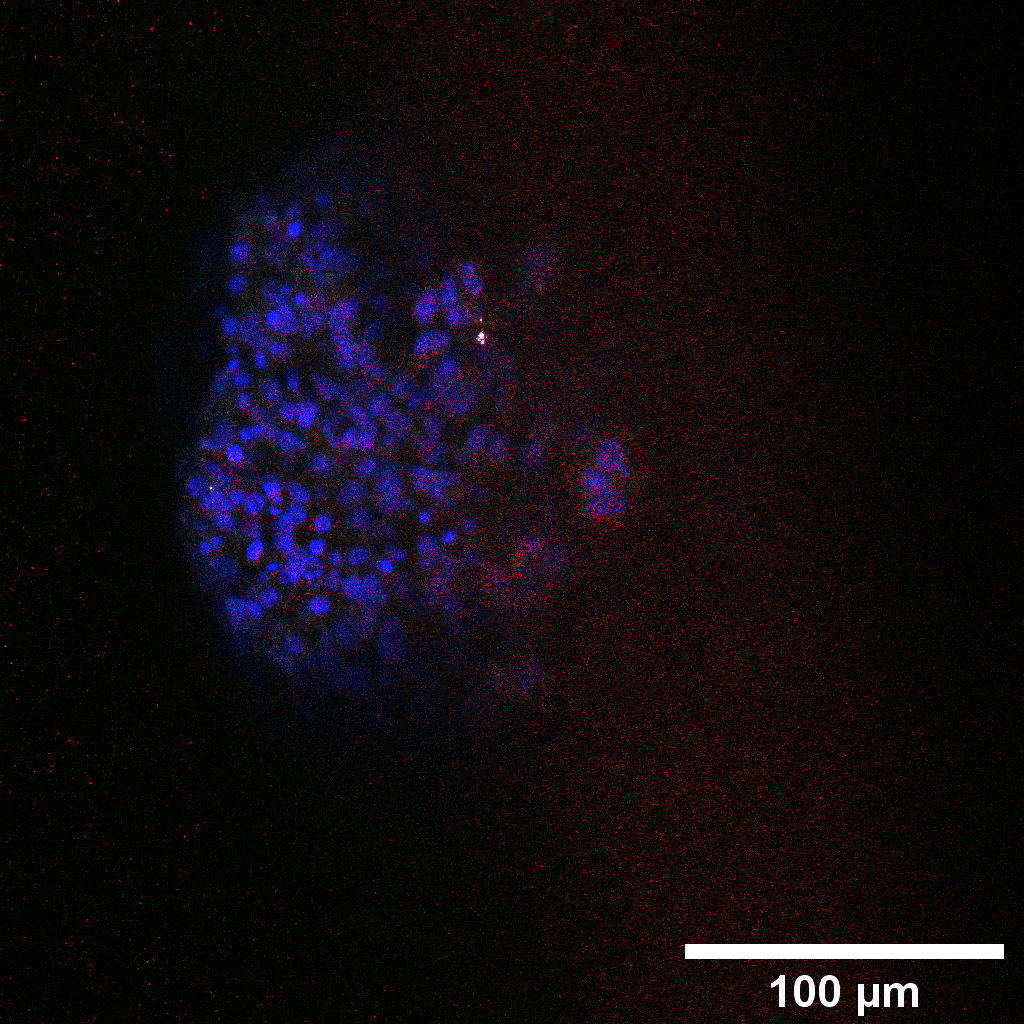
