## Supplementary info for "3D model of mouse embryonic pancreas and endocrine compartment using stem cell-derived mesoderm and pancreatic progenitors"

**Table S1.** Expression of key genes in the 3D in vitro pancreatic aggregates

| **Gene name** | **Function** | **Comments** |
| --- | --- | --- |
| Dll1 | Notch pathway |  |
| Vim | Mesenchymal and EMT marker |  |
| Igfbp2 | Insulin-like growth factor binding protein 2, EMT inducer in pancreatic ductal adenocarcinoma ^1^. Expressed in mesothelial cells in E12.5 and E17.5 mouse pancreas ^2^ |  |
| Irx1 | Differentiation regulators of alpha cells ^2^ |  |
| Yap1 | Hippo pathway |  |
| Epcam | Pancreatic epithelium ^2^ |  |
| Rbpj | Notch pathway |  |
| Id2 | Id DNA-binding protein family. Expressed in mesenchymal population in the mouse embryo pancreas ^2^ |  |
| Hes1, Hes5 | Notch pathway |  |
| Ptn | Heparin-binding cytokine, expressed in interepithelial and delaminating Ngn3 + cells (EMT) in E14.5 mouse pancreas ^3^ |  |
| Egr1 | Expressed in endocrine progenitors and beta cells in E16.5 mouse embryo ^3^ |  |
| Cxcl12 | Chemokine expressed in mesenchymal cells in E14.5 mouse pancreas ^2^ |  |
| Twist1 | Delamination and migration markers |  |
| Col3a1 | Mesenchymal cells ^2^ |  |
| Pdgfrb | Mesenchymal cells in the mouse embryo pancreas ^4^ and expressed in pericytes, which support the formation and maturation of blood vessels ^5^. |  |
| Tcf4 | Transcription factor in Wnt pathway |  |
| Barx1 | Mesenchymal cells | 228 cells: PP_org (18 cells), PP+Matrigel (72 cells), PP+MP_B1 (34 cells) and PP+MP_B2 (104 cells) |
| Bmp7 | Exocrine progenitors | 1415 cells: PP_org (267 cells), PP+Matrigel (857 cells), PP+MP_B1 (41 cells) and PP+MP_B2 (250 cells) |
| Col1a1 | Mesenchymal cells in E14.5 mouse pancreas ^2^ |  |
| Spp1 | Ductal and proliferating ductal cells in E14.5 mouse pancreas ^2^ |  |
| Ptf1a | Exocrine progenitors |  |
| Mecp2 | Methyl-binding protein, expressed in beta cells ^6,7^ |  |
| Atf2 | Beta cells ^2^ |  |
| Gcg | Alpha cells | 0 cells |
| Ins1, Ins2 | Beta cells | 0 cells |
| Tph1 | Beta cells | 9 cells: PP+Matrigel (3 cells), and PP+MP_B2 (6 cells) |
| Sst | Delta cells | 286 cells: PP_org (31 cells), PP+Matrigel (43 cells), PP+MP_B1 (36 cells) and PP+MP_B2 (176 cells) |
| Ghrl | Epsilon cells | 8 cells: PP+Matrigel (3 cells), and PP+MP_B2 (5 cells) |
| Cdh2 (Ncad) | Cell adhesion molecule, required to establish the dorsal pancreatic mesenchyme ^8^. Plays a role in cell-to-cell junction formation between pancreatic beta cells and neural crest stem cells. Expressed in mesoendodermal cells ^9^ |  |
| Onecut1 | Pancreatic progenitors |  |
| Pecam1 | Endothelial cells | 129 cells: PP_org (40 cells), PP+Matrigel (34 cells), PP+MP_B1 (7 cells) and PP+MP_B2 (48 cells) |
| Erg | Endothelial cells | 6 cells: PP+MP_B2 (6 cells) |
| Sox4 | Early endocrine progenitors ^3^ |  |
| Insm1 | Endocrine progenitors. Suppresses Neurod1 and insulin-secreting cells |  |
| Myt1 | Endocrine progenitors |  |
| Stmn2, Stmn3 | Regulator of microtubule stability. Stmn2 is expressed in alpha cells. |  |
| Chga, Chgb | Endocrine cells |  |
| Peg10 | Mainly alpha but also beta cells ^2^ |  |
| Hoxb4, Hoxd4 | Anterior hox genes |  |
| Hoxb6, Hoxc6 | Central hox genes |  |
| Hoxc10 | Posterior hox gene |  |
| Gng12 | Mainly beta, bet also alpha ^2^ |  |
| Itga6 | Exocrine cells: epithelial duct cells. |  |
| Muc1 | Ductal cells | 57 cells: PP_org (15 cells), PP+Matrigel (21 cells), PP+MP_B1 (9 cells) and PP+MP_B2 (12 cells) |
| Rbp1 | Cellular retinol-binding protein type I, |  |
| Pax6 | Pancreatic progenitors, endocrine progenitors, and beta cells. |  |
| Crabp1 | Cellular retinoic acid binding protein I, expressed in endocrine and exocrine glands. |  |
| Cyp26b1 | Involved in the metabolism of retinoic acid. Plays a role in pancreas development. |  |
| Top2a | Proliferating acinar and ductal cells |  |
| Foxm1 | Beta cells |  |
| Ccnd2, Ccnd1, Ccnb2, Ccna2 | Cyclin genes, proliferating cells and endocrine cells ^2,10–12^ |  |

**Table S2.** Marker genes of the 20 clusters of the integrated in vitro-embryo data (Excel spreadsheet)

**Table S3.** List of genes with MI ≥0.23 (Excel spreadsheet)

**Table S4.** Endothelial compartment in each sample

|  | **E12** | **E14.B1** | **E14.B2** | **E17** | **PP_org** | **PP+Matrigel** | **PP+MP_B1** | **PP+MP_B2** |
| --- | --- | --- | --- | --- | --- | --- | --- | --- |
| Endothelial cells out of total sample in percentage | 0.63% | 0.92% | 1.04% | 4.6% | 0.24% | 0.46% | 0.62% | 1.55% |

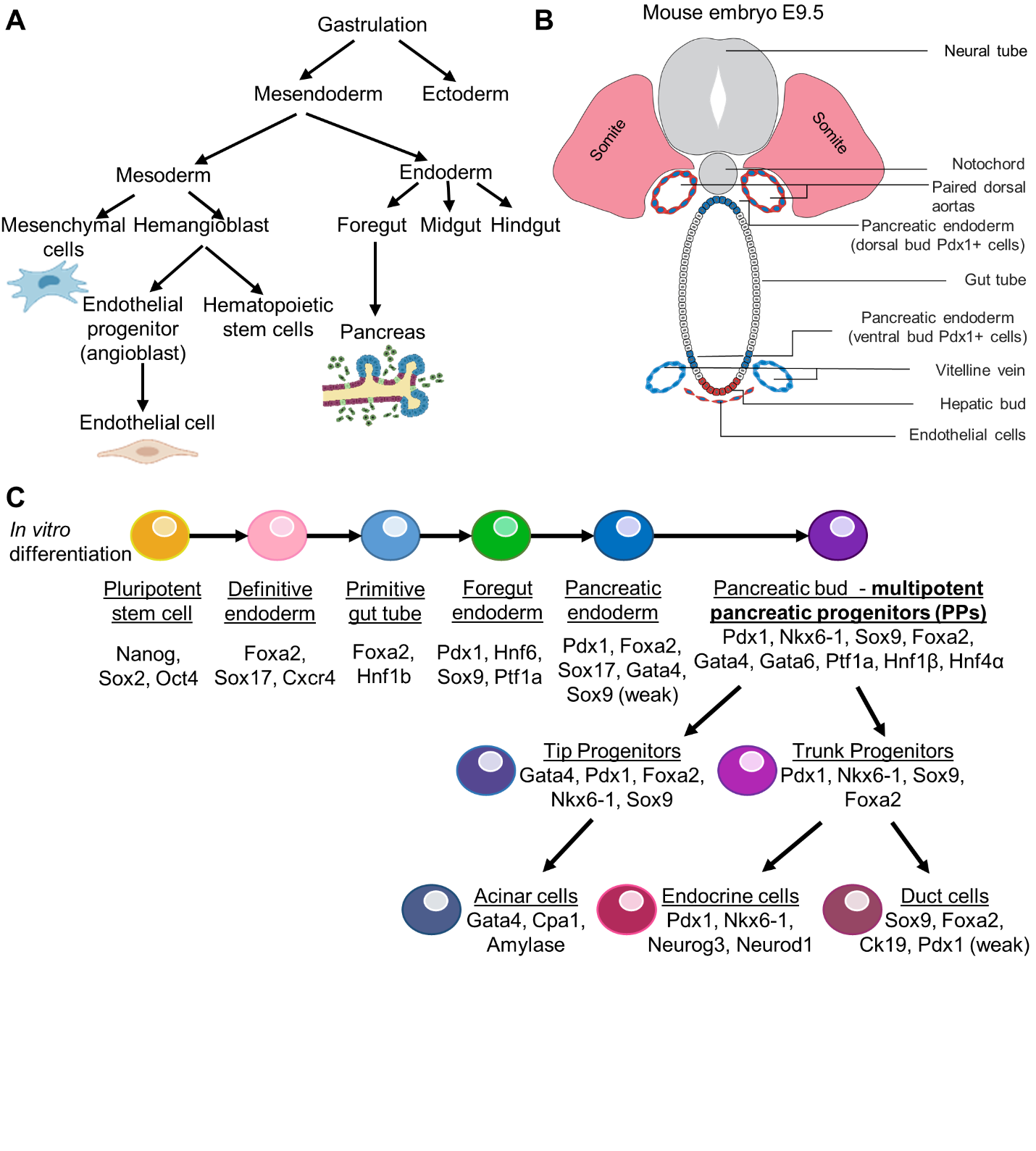

**Figure S1. Pancreas development in the embryo and in vitro differentiation of pluripotent stem cells to pancreatic cell types. A.** Mouse embryo cell differentiation after gastrulation**.** The pancreas is derived from the foregut which is of endoderm origin. Endothelial and mesenchymal cells derive from the mesoderm germ layer. **B.** Cellular interactions in a E9.5 mouse embryo. On E9.5, pancreatic bud formation is induced by the signals secreted in proximal regions. Vascular endothelial growth factor (VEGF) is secreted by the neural tube and the foregut, and triggers dorsal aorta development. The dorsal aorta is composed only of endothelial cells that are in contact with the foregut endoderm. These endothelial cells instruct the formation of the pancreatic bud. This illustration was adapted with some modifications from ^5^. **C.** Suggested in vitro pluripotent stem cell differentiation pipeline and marker genes in each state summarised from studies on murine pancreas development and new insights into human pluripotent stem cells differentiation towards pancreatic identity ^2,9,13,14^.

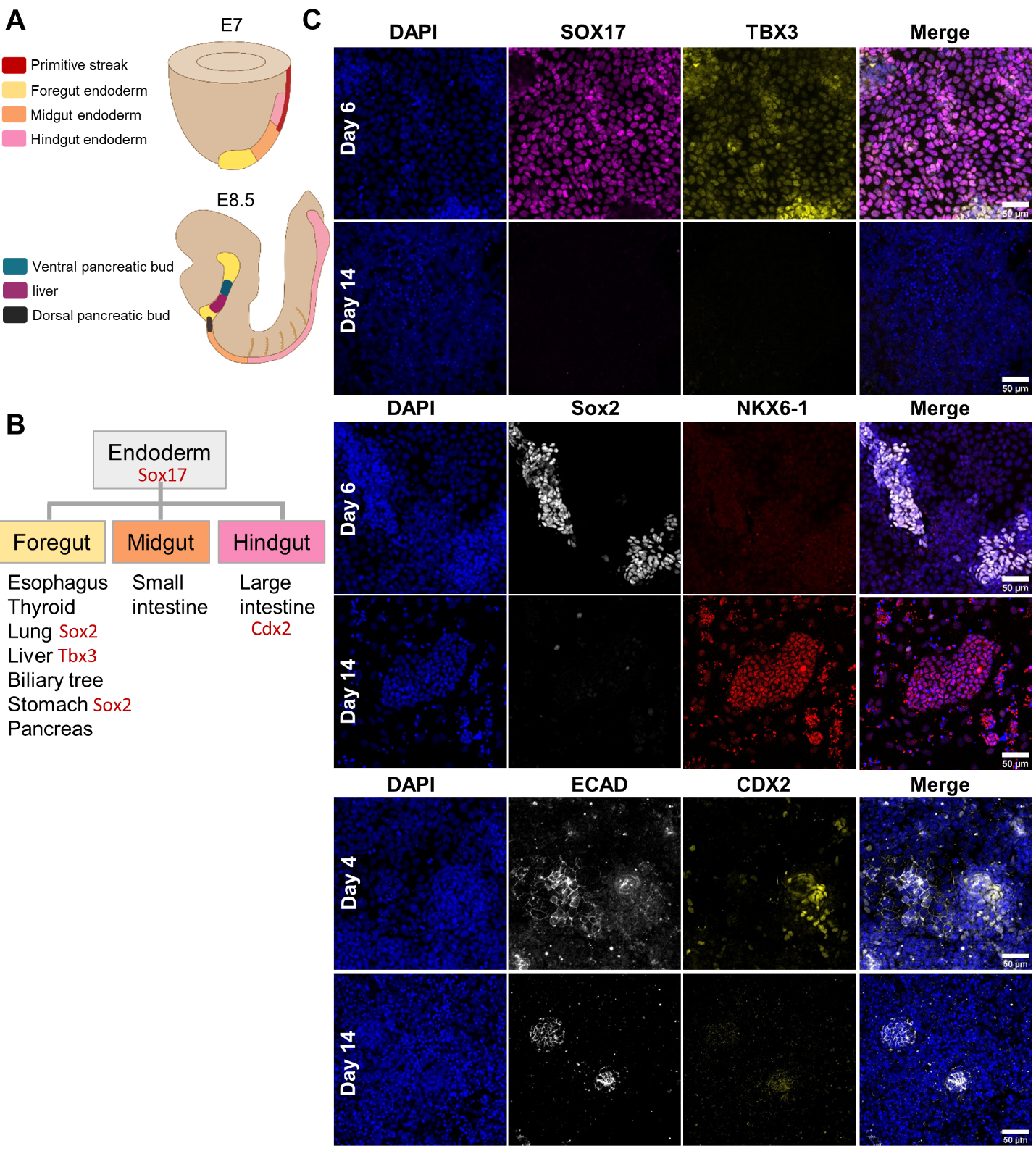

**Figure S2. EpiSC differentiation to pancreatic progenitors is specific. A.** Illustration of endoderm specification in the mouse embryo at E7 and E8.5. **B.** Endodermal organ development and marker genes. **C.** Confocal images along the course of EpiSC differentiation on day 4 and day 6, reflecting the definitive endoderm or gut tube endoderm stages, respectively, and on day 14, i.e., the end of differentiation. The cells were immunofluorescently stained for the markers listed in B, and for the pancreatic marker NKX6-1 and the epithelial marker ECAD. Note that at the end of differentiation, expression of the non-pancreatic endodermal markers diminished.

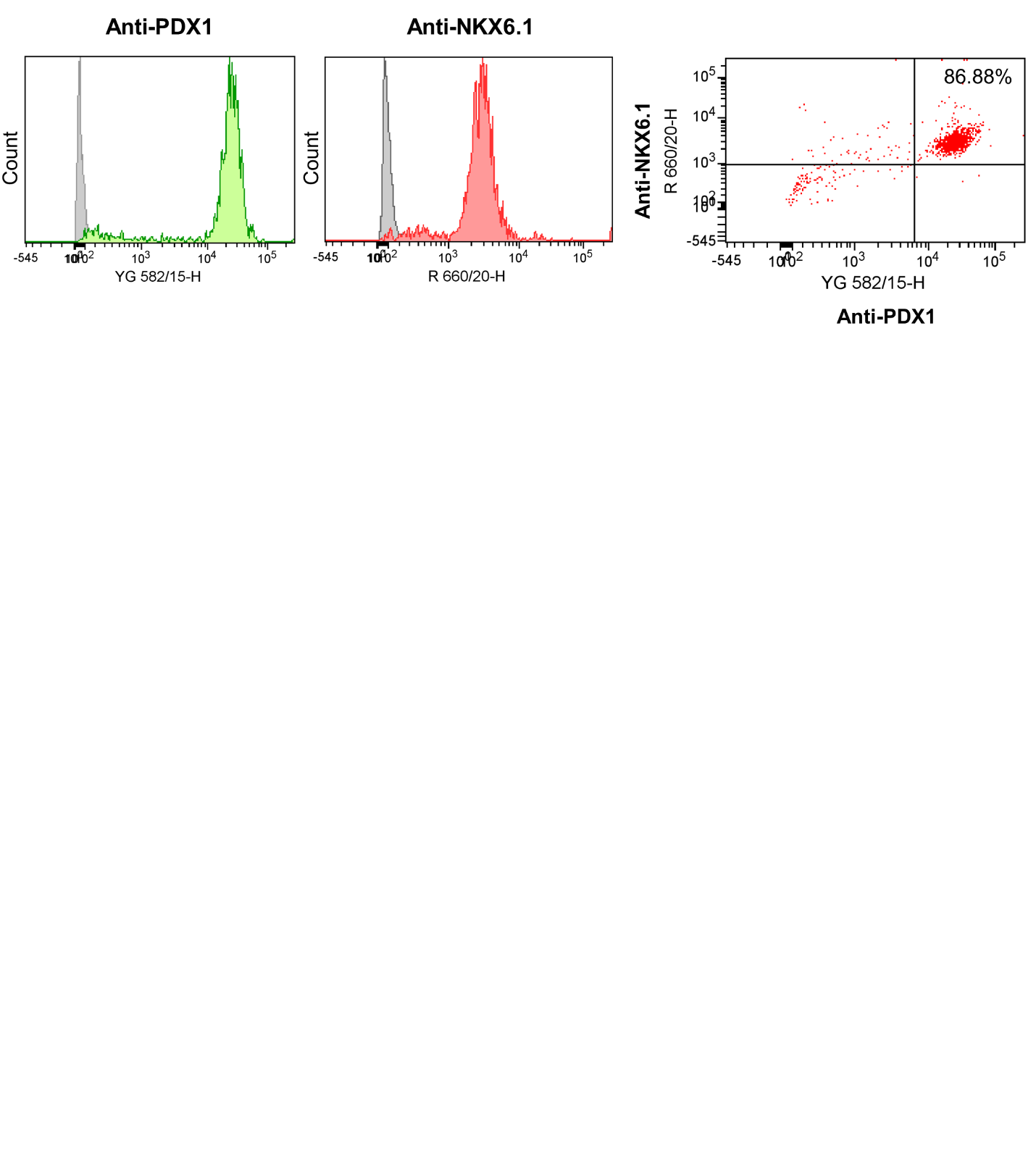

**Figure S3. High yield of PDX1/NKX6-1 double-positive cells on day 14 of EpiSC differentiation to PPs**. Flow cytometry quantification of PDX1/NKX6-1 double-positive cells on day 14 of EpiSC differentiation to PPs (n=2 biological experiments, see Figure 2C).

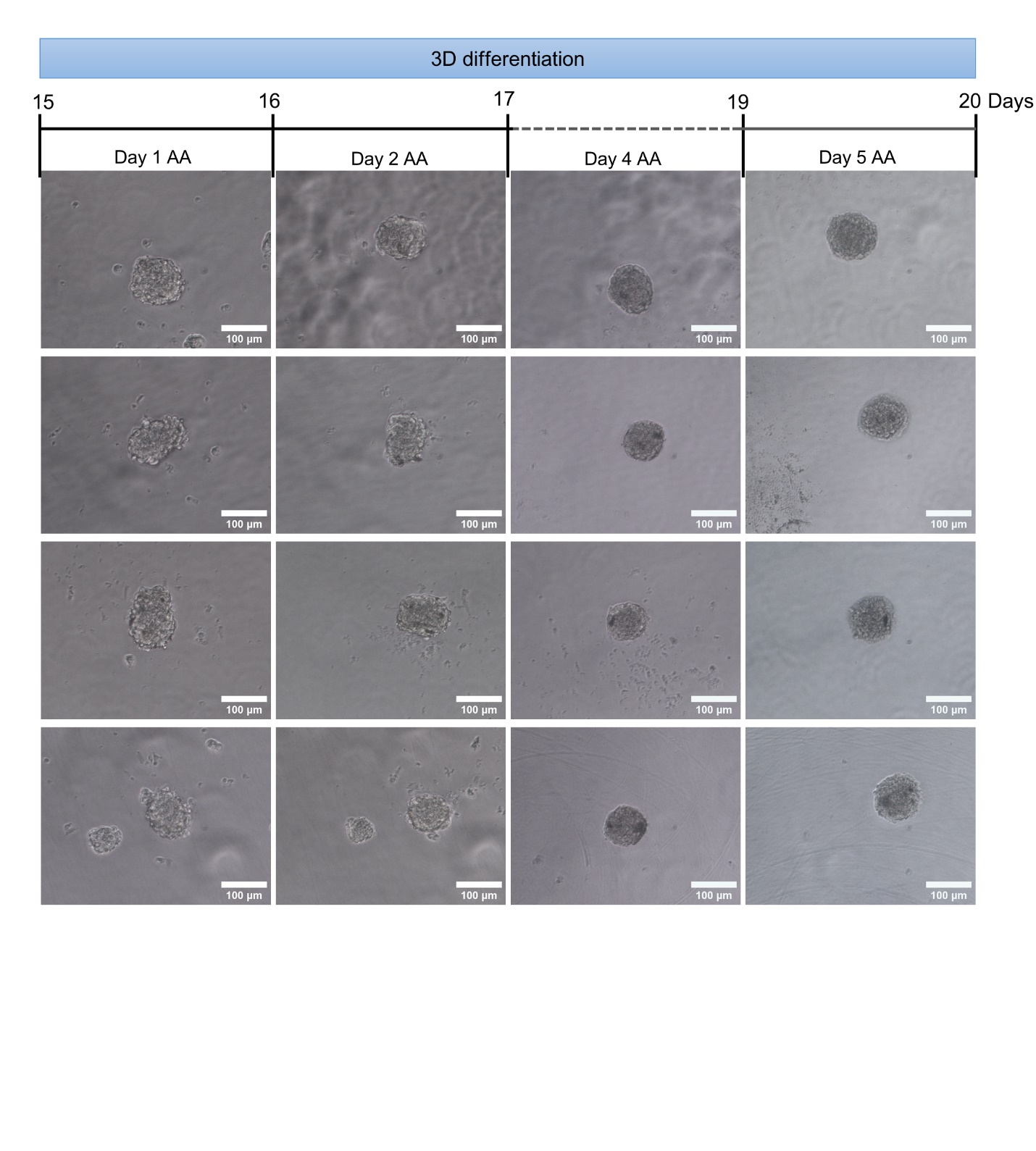

**Figure S4. Aggregates do not grow without mesoderm progenitors.** On day 14, 1000 PPs were plated in each well of a U-bottom 96-well plate. Brightfield images of the aggregates on days 1, 2, 4 and 5 after aggregation (AA). No change in the aggregate size was observed over time.

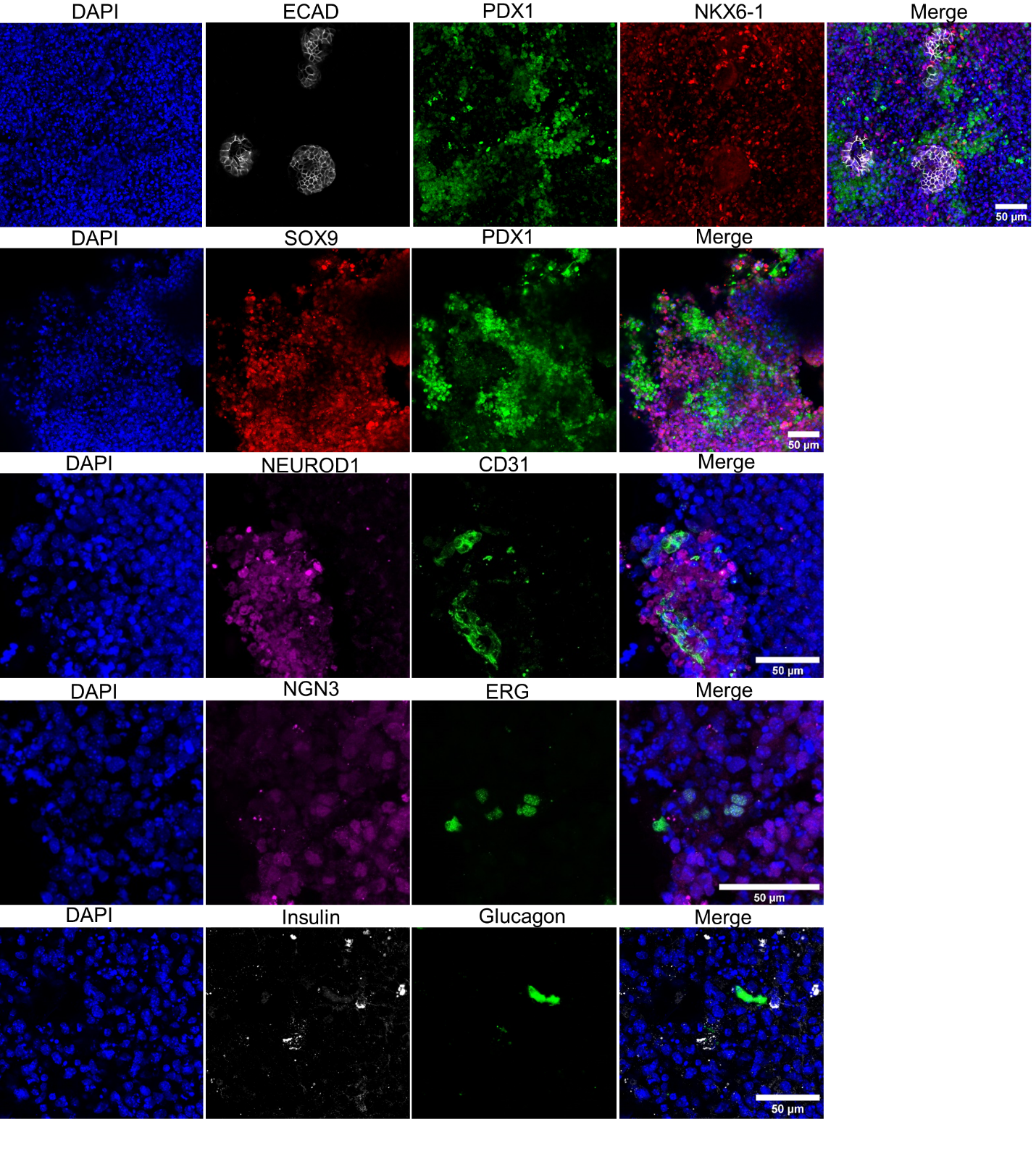

**Figure S5. 3D aggregation of EpiSC-derived pancreas and mesoderm progenitors resulted in the expression of various pancreas cell types.** Confocal images of the aggregates on day 8 after aggregation immunoflourescently stained for the progenitor markers NKX6-1, PDX1 and SOX9, endocrine markers NEUROD1 and NGN3 (Neurog3), endothelial markers ERG and CD31, an acinar marker AMY (amylase), an epithelial marker ECAD, beta cell marker insulin and alpha cell marker glucagon.

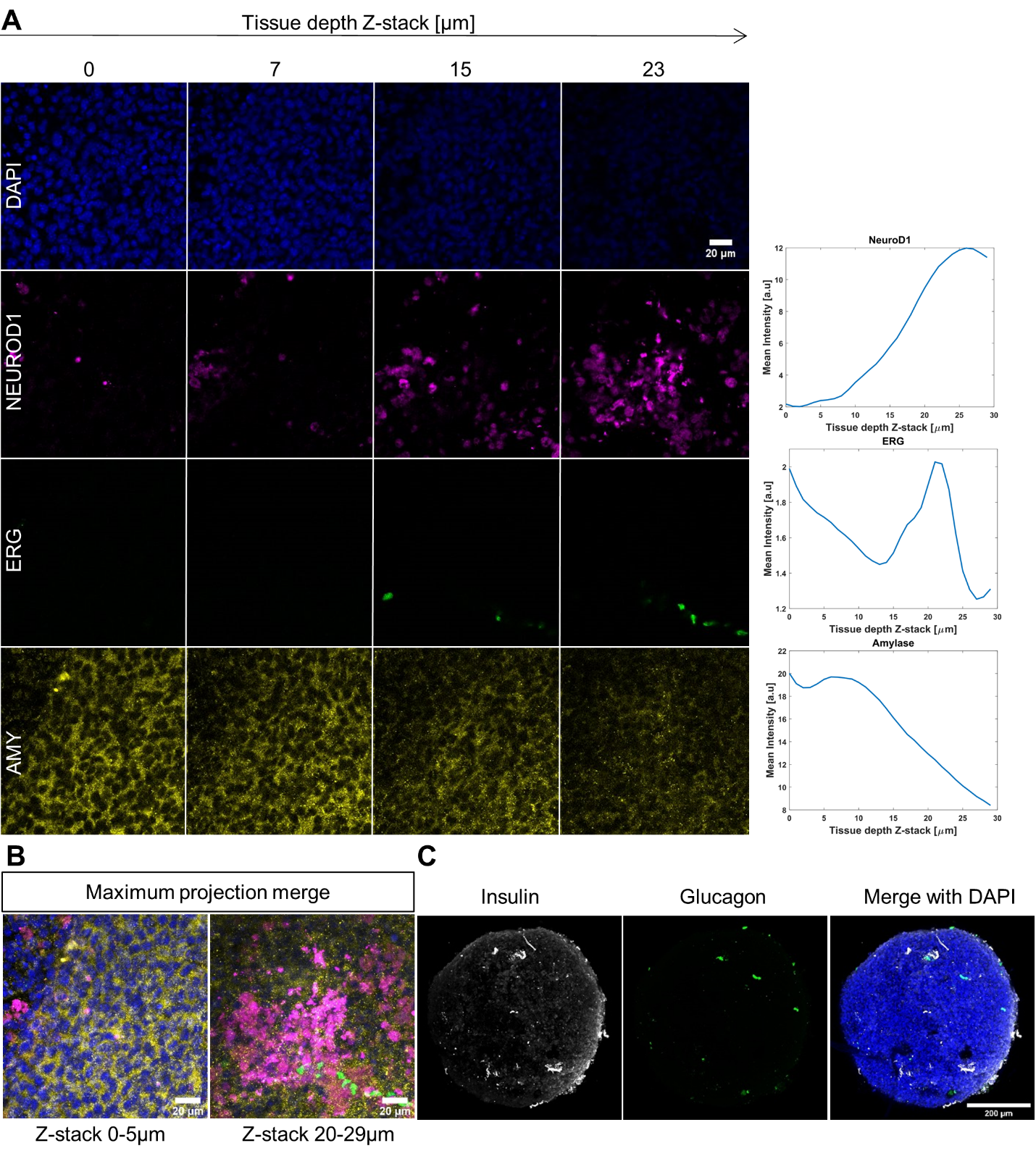

**Figure S6. Endocrine cells are in proximity to endothelial cells. A.** Left: confocal images of the PP+MP aggregates on day 8 after aggregation, immunofluorescently stained for NEUROD1, ERG and AMY along the aggregate depth (Z-stack). Right: The average intensity of each gene along the aggregate depth. **B.** Maximum fluorescence intensity projection of the superficial Z-stacks (0-5µm) and inner Z-stacks (20-29µm) of the aggregate confocal images as in A. **C.** Confocal images of a pancreatic aggregate on day 8 AA immunofluorescently stained for glucagon and insulin, which mark alpha and beta cells, respectively. Nuclear staining with DAPI is in blue.

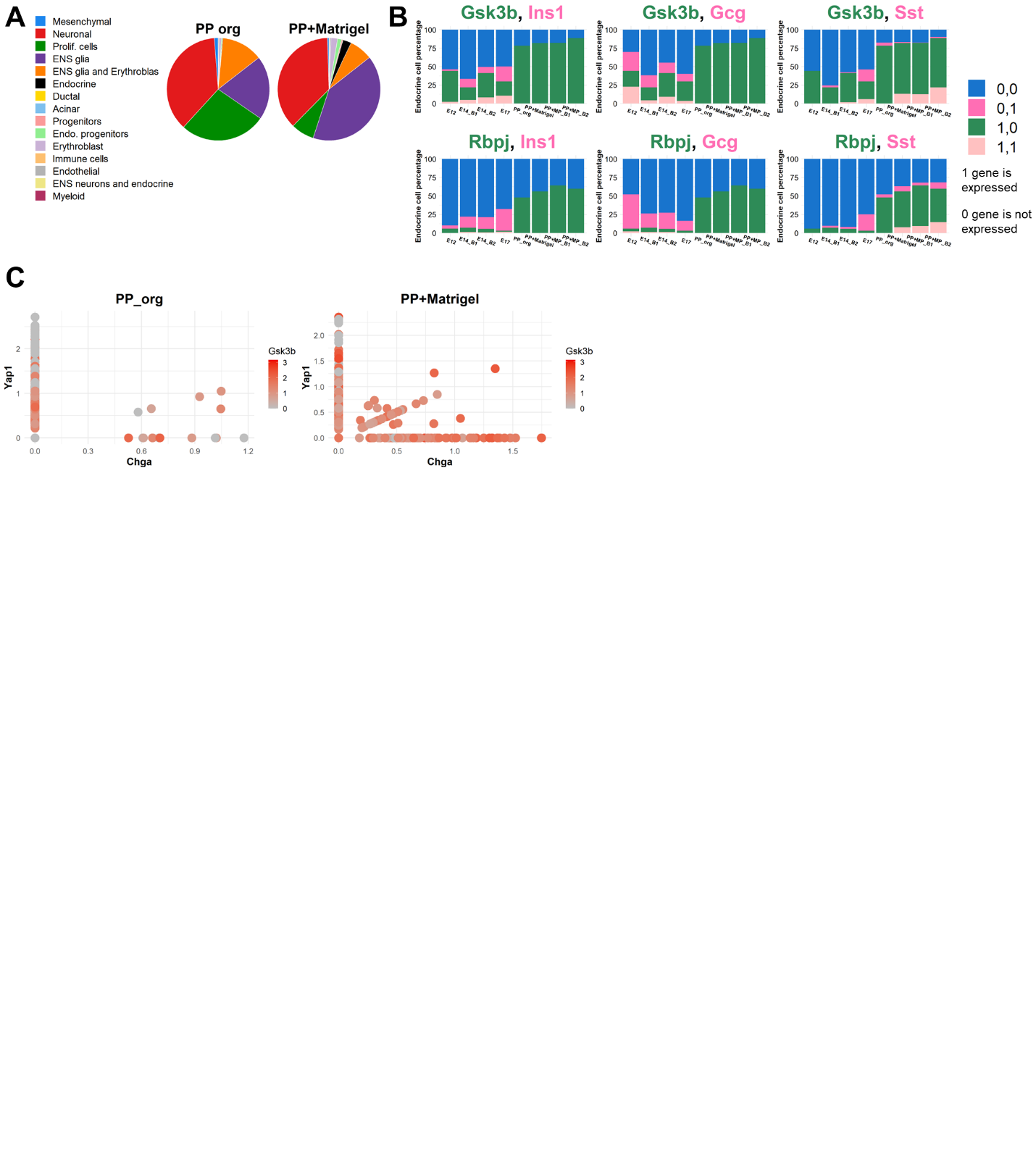

**Figure S7. PP aggregates and PP mixed in Matrigel mainly have a neuronal identity. A.** Pie plots showing the proportion of the different cell types in the PP_org and PP+Matrigel samples. **B.** Endocrine cell composition (clusters: endo. progenitors, endocrine and ENS neurons and endocrine) of cells expression either Gsk3b or Rbpj and the hormones Ins1, Gcg or Sst. 0,0 indicates expression of neither the two genes in a cell, 0,1 indicates no expression of Gsk3b or Rbpj and expression of the hormones (Ins1, Gcg or Sst) in a cell, 1,0 indicates expression of Gsk3b or Rbpj and no expression of the hormones (Ins1, Gcg or Sst) in a cell and 1,1 indicates double expression of the two genes in a cell. **C.** Gene expression of Chga, Yap1 and Gsk3b in PP_org and PP+Matrigel samples. Each dot represents a cell, x and y axes show the normalized expression of Chga and Yap1, respectively. The normalized expression level of Gsk3b is indicated by the colour intensity.

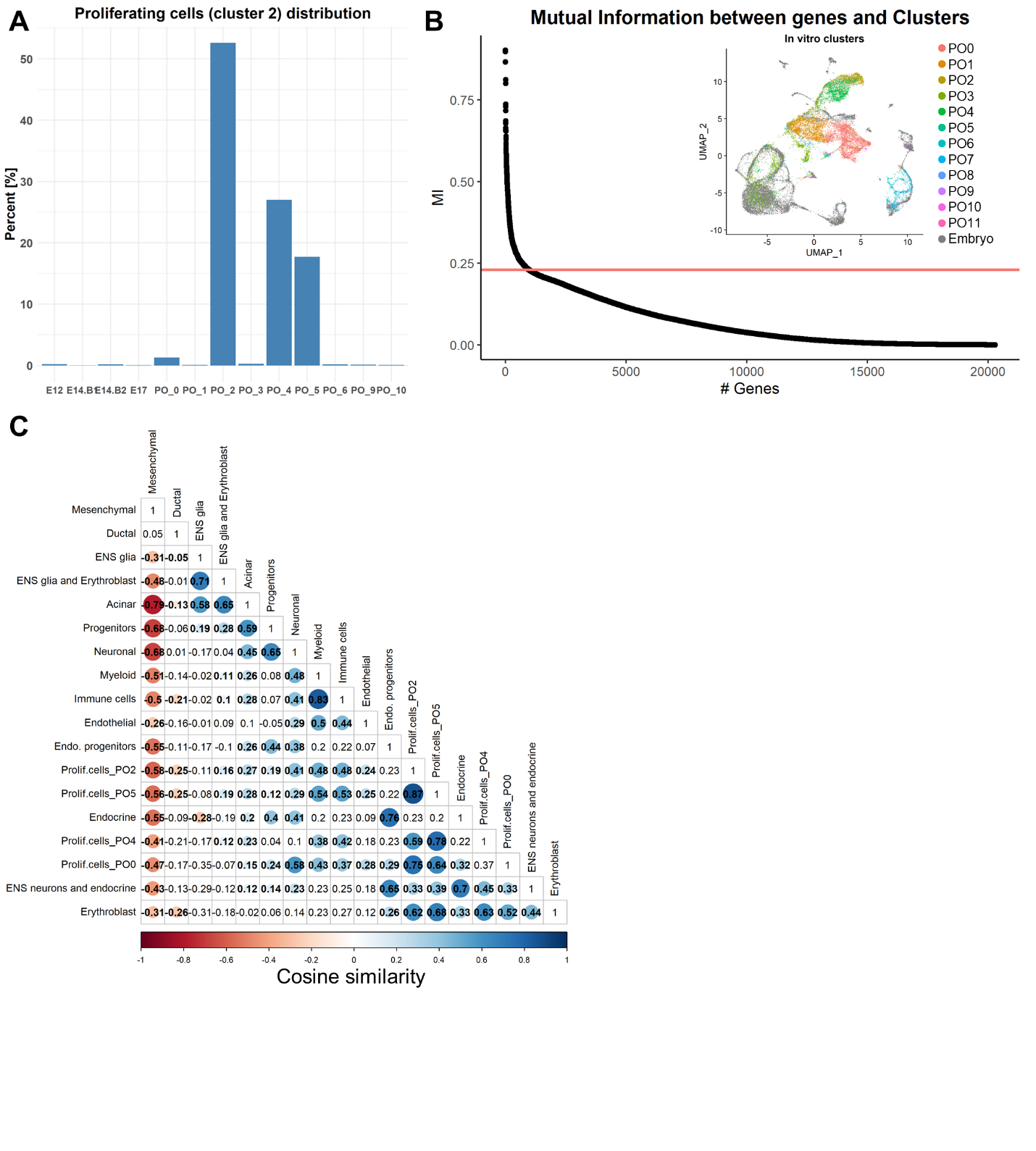

**Figure S8. The major cell population in the 3D in vitro pancreatic aggregate is similar to embryonic erythroblast cells. A.** Composition of the proliferating cell cluster (cluster 2). PO_0 stands for pancreatic aggregate cluster 0, PO_1 stands for pancreatic aggregate cluster 1, and so forth from the in vitro sample clustering (Fig. 4A). **B.** Mutual information (MI) calculation between expression of all the genes and cell assignment to all embryonic clusters. The horizontal red line indicates MI=0.23. Only genes with MI≥0.23 were used to calculate the cosine similarity between the four in vitro clusters in the proliferating cell group (A) and the known embryonic clusters. Presented on the right: projection of the 12 pancreatic in vitro clusters on the integrated UMAP plot. **C.** Cosine similarity score between the four major in vitro clusters comprising the proliferating cell group and known embryonic cell types from the integrated data. Non-blanked cells in the cosine similarity matrix indicate empirical p-value <0.01, colour and size indicate the correlation strength.

**Video S1.** Movie showing confocal image Z-stacks of a total 70.15 µm tissue depth of a PP+MP aggregate embedded in 6.6% gelatin crosslinked with 5% microbial transglutaminase (mTG). CD31 (red) marks the vessel-like network and DAPI (blue) marks the nuclei.
